# Ecological tristability driven by total carbon availability over resource complexity in a synthetic microbial community

**DOI:** 10.64898/2026.02.26.708221

**Authors:** Anna M Bischofberger, Johannes Cairns, Inga-Katariina Aapalampi, Sanna Pausio, Meri Lindqvist, Ville Mustonen, Teppo Hiltunen

## Abstract

Even though complex microbial communities are ubiquitous and provide essential services for natural and human-associated ecosystems, our knowledge about their assembly and dynamics is incomplete. There is an ongoing debate whether the behavior of complex communities can be predicted from the outcome of pairwise competition of species, and whether communities reach alternative stable states depending on the level and complexity of resource provided for growth. To estimate the effect of two resource gradients, total carbon availability and resource complexity, on the compositional dynamics of a complex microbial community, we conducted a 16-day serial passage experiment, transferring a 16-species synthetic community in 96 different resource environments. We observed that although both resource dimensions influenced community composition, total carbon exerted a considerably larger effect. Additionally, we saw the emergence of a tristable pattern along the total carbon gradient, a feature not observed for the resource complexity gradient. Using monoculture assays, we identified lag phase duration as the dominant predictor of competitive success at carbon extremes, with maximum growth rate increasing in importance as lag times converged. Total carbon availability thus structured community state transitions and regulated which growth trait governed competitive sorting. These results suggest the importance of total carbon level over resource complexity and identifying dominant species for the quest to successfully manage, maintain and manipulate complex microbial communities.

## Introduction

Microorganisms are part of our surroundings, where they exist in diverse, often complex communities. We depend on the functioning of these communities not only for the services they provide in natural ecosystems ^1–3^ but also for maintaining human health ^4^ and a broad variety of applications such as food production, wastewater treatment, and environmental bioremediation ^5–7^. Identifying the mechanisms that govern microbial community assembly and dynamics and the conditions under which they are valid is thus crucial. However, although high-throughput sequencing has revealed immense microbial diversity across environments, the processes determining species co-existence, dominance, and exclusion are still incompletely understood and hence remain a central challenge in microbial ecology.

Classical ecological theory predicts that community diversity is constrained by resource availability. The competitive exclusion principle states that the number of species able to co-exist is limited by the amount of resources, implying that increases in resource quantity or diversity should promote species richness ^8,9^. In microbial systems, empirical work has supported this framework by showing that higher nutrient supply can relax competitive constraints and increase diversity. However, microbial communities frequently fail to match predictions based on simple resource competition models, suggesting that additional mechanisms facilitate co-existence ^10,11^.

Metabolic cross-feeding is one such mechanism. By converting primary substrates into secondary metabolites that can be exploited by other taxa, cross-feeding effectively expands niche space even when communities are supplied with a single resource ^12,13^. Experimental studies have demonstrated that such metabolic interactions can generate stable multispecies communities and reproducible assembly patterns that cannot be explained by resource partitioning alone ^14^.

Beyond resource quantity, the metabolic complexity of available substrates can strongly influence community structure. Silverstein *et al.* (2024) ^15^ have proposed the divergence-complexity hypothesis which adheres to the two principles of ‘metabolic diversity begets [microbial] diversity’ and ‘[metabolic] diversity begets divergence [of microbial communities]’. By culturing soil microbial samples with differing initial community composition at t_0_ in increasingly complex media formulations, they showed how metabolically complex resources provide a greater diversity of available niches, leading to increased divergence in community composition. Other studies have provided results in accordance with this hypothesis, both in natural and synthetic communities ^16,17^. Faced with ever greater disruption of natural environments due to human activity, it is essential to further our understanding of the dynamics behind regime shifts in and the resilience of microbial communities ^18^.

The search for further factors promoting co-existence in microbial communities leads to differences in life-history traits. Though known to modulate microbial competition, they are often underintegrated into community assembly frameworks. In particular, differences in maximal growth rates and/or lag-phase duration can critically influence competitive outcomes. The lag phase, a shift in metabolism from fermentation to respiration^19^, reflects active physiological adjustment to novel environments and varies among species, environmental conditions, and even among individual cells ^20–22^. Using donor stool communities and reconstituted synthetic versions of those communities, Aranda-Dìaz *et al.* showed that removing the fastest growing, intially dominant strains results in the second fastest strain reaching the highest abundance.^23^ The most successful strains exhibited a combination of high growth rate and short lag phase. Others have provided evidence, that in batch and serial-transfer systems, shorter lag phases alone can confer priority effects, allowing for early resource capture that may outweigh differences in maximal growth rate ^24,25^. Experimental evidence indicates that such priority effects can generate historical contingency and alternative community states ^26^. However, growing evidence suggest that the outcome of such community culturing depends not only on the properties of the participating species but also on a range of environmental and/or procedural factors.^27–30^

Environmental stress further shapes the balance between competition and facilitation. The stress-gradient hypothesis (SGH), originally developed in plant ecology, predicts that competition dominates under benign conditions, whereas facilitative interactions become more important under increasing stress, often resulting in hump-shaped diversity patterns ^31,32^: at high environmental stress, the conditions are so harsh that few but the hardiest and best adapted species can survive; when environmental stressors are all but absent, organisms are able to invest resources in their competitive abilities, driving less-competitive species into extinction. Recent studies suggest that analogous dynamics operate in microbial communities, where nutrient limitation or other stressors such as heavy metal contamination, water availability or elevation can shift interactions from competition toward facilitation ^33–37^.

Though recent advances in sequencing have helped elucidate the high diversity of microbial communities in different conditions and the availability of high-power computing allows for the quick processing of large datasets^38–40^, there are still gaps in our ability to predict long-term community outcomes in changing environments. While there is evidence from some experiments that the complexity of community dynamics can be reduced to pairwise competition of species^41^, there are other studies that highlight the importance of higher-order interactions for the development of multispecies communities^42^.

Here, we hypothesised that increasing resource complexity, implemented as a gradient of metabolically simple to complex media (R2A-based), would increase diversity in a multispecies synthetic microbial community. Using a fully factorial design of 96 resource environments, we exposed the community to two orthogonal nutrient gradients: total carbon availability and increasing metabolic complexity. Contrary to expectations, we found that total carbon availability was the primary driver of community-level regime shifts, with resource complexity exerting a secondary, modulatory effect, and that the emerging tristable pattern could be predicted from monoculture growth traits of the dominant species.

## Materials & Methods

### Synthetic bacterial community

In this study, we used a modified, 16-species version of our synthetic community that has been used in various earlier studies ^43–45^. All species were obtained from the HAMBI culture collection (University of Helsinki) and represent diverse taxonomic groups and ecological origins, including aquatic, soil, and host-associated environments. The community consisted of the following species: *Pseudomonas putida* (HAMBI-ID 6), *Agrobacterium tumefaciens (*105), *Comamonas testosteroni* (403), *Citrobacter koseri* (1287), *Morganella morganii* (1292), *Kluyvera intermedia* (1299), *Sphingobium yanoikuyae (1842), Sphingobacterium spiritivorum* (1896), *Aeromonas caviae* (1972), *Pseudomonas chlororaphis* (1977), *Bordetella avium* (2160), *Cupriavidus oxalaticus* (2164), *Paracoccus denitrificans* (2443), *Stenotrophomonas maltophilia* (2659), *Niabella yanshanensis* (3031), and *Microvirga lotononidis* (3237).

### Media preparation and composition

For the serial passage experiment (Fig. 01), we prepared a total of 96 distinct growth media that varied along two experimental gradients: total carbon concentration and resource complexity (% R2A carbon contribution).

**Figure 01.**
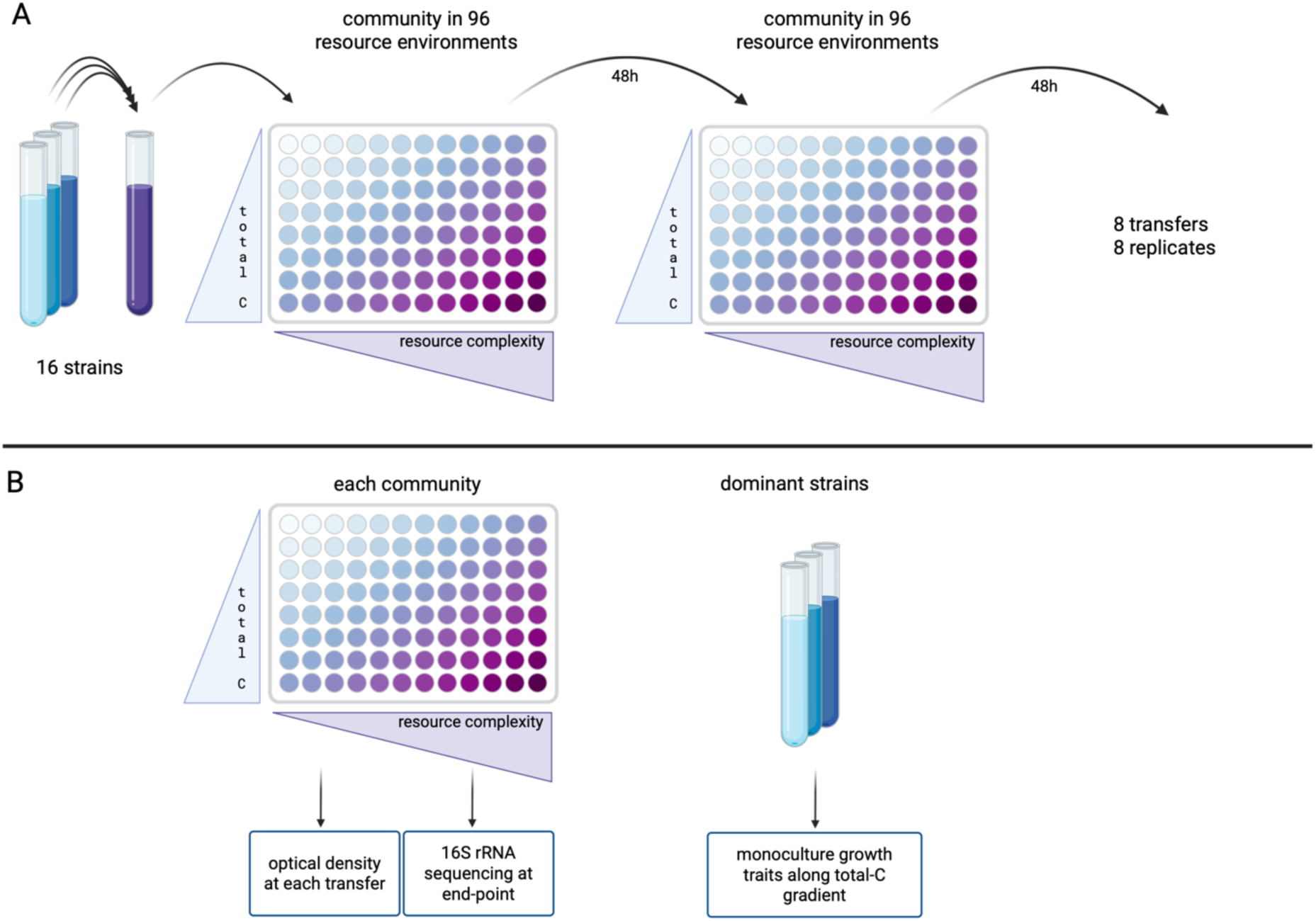
Schematic representation of experimental setup. A | After pre-culturing each of the 16 species of the community for one growth cycle, we adjusted optical density of each culture, pooled them at equal volume and inoculated 96 resource environments, with each condition replicated 8 times. Each culture was serially transferred for 16 days, with 1% of the culture transferred every 48 h. B | At each transfer, we measured OD_600_ of each community. In addition, we conducted 16S rRNA sequencing of all communities at the end-point and measured growth traits of each of the dominant species (*A. caviae, P. chlororaphis, C. koseri*) in monoculture in all carbon levels in the absence of R2A. Alt text. Two-panel schematic representing the study set-up. Panel A describes parameters of the serial passaging experiment. Panel B explains analytical procedures of the community and single-species experiments.

First, a glucose stock solution was prepared by dissolving glucose in deionized water and filter-sterilized. M9 minimal medium was prepared according to standard protocols by dissolving 11.28 g of M9 minimal medium salts (MP Biomedicals, UK) in 1.0 L of Milli-Q purified water and supplementing with MgSO₄ and CaCl₂. Glucose minimal media were prepared by adding glucose stock to M9 minimal medium to obtain a maximum carbon concentration of 4 g C L⁻¹. R2A medium was prepared by dissolving R2A Broth powder (Neogen, UK) in M9 minimal medium. The total carbon content of R2A powder was determined at Lammi Biological Station (Finland) using a Shimadzu TOC-L csh analyzer (SFS-EN 1484), showing that R2A contains 272 mg C per gram of dry powder. Based on this measurement, R2A was added to obtain a maximum concentration of 4 g C L⁻¹, corresponding to 14.7 g R2A powder per liter. Glucose minimal medium (4 g C L⁻¹) and R2A medium (4 g C L⁻¹) were mixed in defined proportions to generate 12 media in which the proportion of R2A-derived carbon ranged from 0 to 100 % of total carbon (resource complexity gradient), while keeping total carbon constant at 4 g C L⁻¹. Each of these 12 media was then diluted 1:2 with carbon-free M9 minimal medium in a serial dilution series to generate eight total carbon concentrations ranging from 0.031 to 4 g C L⁻¹. This resulted in a total of 96 distinct media compositions, which are listed in Table S1.

For each condition, 1000 µL of medium was dispensed into 96-well deep-well plates (Sarstedt, Germany). Each of the 96 growth media compositions was replicated eight times.

### Pre-culture and community assembly

All 16 bacterial species were stored at -80 °C in glycerol stocks. Each species was grown separately from frozen stocks in 25mL microcosm flasks containing 6 mL of nutrient-rich proteose peptone yeast extract medium (PPY; 20 g proteose peptone and 2.5 g yeast extract per 1 L of deionized water). Cultures were incubated for 48 h at 30 °C with constant shaking at 70 rpm. After incubation, cultures were washed twice with unsupplemented M9 medium and the optical density at 600nm (OD_600_) was adjusted to 1.4 using unsupplemented M9 medium. Equal volumes of each adjusted culture were pooled to form the synthetic community. One milliliter of the pooled community was stored at -20 °C as a baseline control for amplicon sequencing.

### Serial passage experiment

We used serial transfer and culture protocols similar to those used in earlier studies^43,44^ in a serial passage experiment conducted over 16 days and consisting of eight growth cycles. Every 48 h, 1 % of each culture from the previous plates was transferred into new 96-well deep-well plates containing 1 mL of fresh growth medium with the same R2A and glucose-derived carbon concentrations. At each transfer, OD₆₀₀ was measured to monitor bacterial growth. After measurement, samples were frozen at -20 °C for subsequent amplicon sequencing.

### 16S rRNA gene amplicon library preparation and sequencing

Microbial community composition was characterized using 16S rRNA gene amplicon sequencing with a two-step PCR and dual-indexing strategy. Amplicon libraries were prepared using an in-house protocol (see Supplementary methods, primers are listed in Supplementary Tables S12 & S13), based on the TaggiMatrix (Adapterama II) method of Glenn *et al.* (2019)^46^.

Relative abundances of the 16 bacterial species were determined by mapping paired-end 16S rRNA amplicon reads following the analysis workflow described by Cairns *et al.* (2025)^43^. Reads were quality-trimmed, merged, filtered, and mapped to reference 16S rRNA gene sequences for each species. Species-specific read counts were normalized according to 16S rRNA gene copy number.

### Growth assays across a glucose-derived carbon gradient

We measured growth of ancestral bacterial strains for the three species that dominated in the community experiment (see Results section) across a glucose -derived carbon gradient, without R2A: *Aeromonas caviae* (1972), *Citrobacter koseri* (1287), and *Pseudomonas chlororaphis* (1977). Bacteria were grown individually in 6 mL PPY medium in 25 mL microcosm flasks at 30 °C with constant shaking at 70 rpm for 48 h. After incubation, cultures were centrifuged and washed twice with an equal volume of unsupplemented M9 medium and subsequently diluted 1:10’000 for inoculation. Sixteen glucose solutions with carbon concentrations ranging from 0.07-2 g C L⁻¹ (each at 80 % of the previous concentration; see Table S2) were prepared in M9 minimal medium. For each condition, 140 µL of medium was dispensed into 96-well plates.

Ten microliters of each diluted bacterial suspension were added to the corresponding wells of 96-well plates, with six replicates per species at each carbon concentration. Plates were sealed with breathable membranes and incubated in LogPhase600 (Agilent BioTek, USA) devices at 30 °C with shaking at 800 rpm. OD_600_ was measured every 10 min for 48 h. Three serial transfers were performed by transferring 10 µL of culture into fresh plates containing the same carbon conditions at each 48 h interval.

### pH measurements in 16-species community

To assess whether pH changes could influence community composition, we measured changes in pH in the 16-species synthetic bacterial community composed of ancestral strains grown in three glucose solutions with carbon contents of 0.03125, 0.25, and 4 g C L⁻¹. Each condition was replicated three times, with uninoculated media included as controls. A glucose stock solution was prepared in deionized water and filter-sterilized. The highest carbon medium (4 g C L⁻¹, corresponding to 10 g glucose L⁻¹) was prepared by dissolving the sterile glucose stock solution in M9 minimal medium supplemented with MgSO₄ and CaCl₂, and lower carbon concentrations were generated by serial dilution (1:16 and 1:8). The 16-species community used in the pH experiment was initially constructed from individual ancestral strains. Each strain was grown individually in 6 mL 100 % PPY medium in 25 mL microcosm flasks at 30 °C with constant shaking at 70 rpm for 48 h. Then, 100 µL of each culture was transferred to 6 mL of 100 % R2A medium and incubated for 24 h at 30 °C. Equal volumes of the R2A pre-cultures were centrifuged and resuspended in unsupplemented M9 medium. The cultures were then combined in equal volumes and mixed 1:1 with 85 % glycerol to generate a frozen stock stored at -80 °C.

For the pH experiment, 50 µL of this stock solution was inoculated into two 25 mL microcosm flasks containing 6 mL of nutrient-rich 100 % R2A medium. Cultures were incubated for 24 h at 30 °C with constant shaking at 70 rpm, and OD₆₀₀ was measured. Next, 5 mL of each pre-culture was centrifuged at 3000 × g for 3 min. The supernatant was removed, and pellets were resuspended in 4 mL of unsupplemented M9 medium before combining the cultures. OD_600_ was measured, and 500 µL of the combined community was inoculated into Erlenmeyer flasks containing 50 mL of M9 minimal medium supplemented with glucose at the three carbon concentrations. Cultures were incubated at 30 °C for 48 h. The pH of each medium was measured immediately after inoculation and again after 48 h of incubation using a calibrated pH meter. In addition, the pH of the glucose stock solution and uninoculated media at each carbon concentration was measured as baseline control values.

### pH measurements for individual species

We found that, in line with our expectation that pH would be lower at the high-end of the carbon gradient due to byproduct secretion, pH was reduced by increasing carbon levels in the community setting but we could not decouple this from OD / microbial productivity effects. We therefore tested individual species. All 16 species used in the serial passaging experiment were grown individually in M9 minimal medium supplemented with glucose at a carbon concentration of 4 g C L⁻¹. Cultures were grown in 10 mL volumes in 25 mL microcosm flasks, and the pH of the medium was measured prior to inoculation from separate flasks. Pre-cultures were prepared by retrieving each species from frozen glycerol stocks and growing them separately in 25 mL microcosm flasks containing 6mL of 100 % R2A medium at 30 °C for 2 h with constant shaking at 70 rpm. OD₆₀₀ was measured both after pre-culture growth and again after washing. One milliliter of each preculture was centrifuged at 3000 × g for 3 min. The supernatant was removed, and the pellets were resuspended in 1mL of unsupplemented M9 medium. Then, 100 µL of the washed cultures was inoculated separately into microcosm vials containing 10 mL of M9 minimal medium supplemented with glucose (4 g C L⁻¹). Cultures were incubated at 30 °C for 48 h with constant shaking at 70 rpm. OD_600_ and pH were measured after incubation. The pH of uninoculated medium was measured as a baseline control.

Overall, we found a similar relationship between microbial productivity and pH in the individual species setting, with some notable exceptions (Fig. S1). Importantly, *A. caviae* lowers pH to a lower level than does *C. koseri*. Based on these observations, pH change does occur along the carbon axis, might affect community structure, and is driven more by general microbial productivity than pH reduction by specific species. In conclusion, we were able to rule out pH as a major factor driving community composition in our experiments.

### Statistical analyses

All statistical analyses were conducted in R version 4.4.2^47^. Alpha and beta diversity metrics across analyses were computed using the vegan package^48^.

#### Community composition

To quantify the effects of resource gradients on community composition, we calculated Bray-Curtis dissimilarities from relative abundance matrices and tested for marginal effects of total-C and resource complexity using permutational multivariate analysis of variance (PERMANOVA; 9’999 permutations), as implemented in the vegan package^48,49^. Total carbon concentration and resource complexity (% R2A carbon contribution) were included as continuous predictors, and marginal sums of squares were used to assess their independent contributions to compositional variance.

Low-dimensional ordination of communities was performed using t-distributed stochastic neighbor embedding (t-SNE) with the Rtsne package on the relative abundance matrix, using PCA-initialized inputs, perplexity = 35, θ = 0.5, and 7’000 iterations^50^. To identify discrete compositional states, density-based clustering was applied to the two-dimensional t-SNE embedding using DBSCAN (eps = 4, minPts = 4; dbscan package), with noise points excluded from cluster assignment. To assess the role of the simple versus complex resource gradients in structuring community states, t-SNE ordinations were visualized globally and faceted by total-C concentration (Fig.03). Community repeatability across replicates was quantified as dispersion around group centroids in ordination space, using the betadisper function in the vegan package, with higher dispersion interpreted as lower compositional repeatability (Fig. S4).

#### Monoculture growth traits

We fit linear models using lag time, maximum growth rate, and area under the curve (AUC) as response variables and species identity, total-C level, and their interaction as explanatory variables, performing ANOVA tests on the models. For pairwise comparisons, we calculated estimated marginal means (EMMs) with the emmeans package^51^.

To test whether monoculture traits predict community dominance across strains, we modeled strain relative abundance as a function of standardized lag time, maximum growth rate, and area under the curve (AUC), including strain identity as a covariate. For carbon-level aware analysis, the dominant species abundance data from the community experiment were linked to monoculture trait data at the nearest-matching glucose levels. Analyses were performed separately for the full carbon gradient and for the intermediate transition zone (0.25-1.0 g L^−1^). We quantified model fit using *R*^2^ and estimated the unique contribution of each trait as the change in *R*^2^ upon its removal from the full model. For comparison, we also calculated marginal *R*^2^ values from univariate regressions of relative abundance on each trait.

#### Biomass and diversity

Community biomass (OD_600_) was analyzed with linear models including total-C concentration, resource complexity, and their interaction. Model terms were evaluated using ANOVA on the fitted models, and variance explained was summarized using model *R*^2^ and an additive-vs-interaction model comparison.

To characterize how diversity responded to the two resource axes, Shannon diversity was quantified in two complementary ways. First, linear models of Shannon diversity included total carbon, resource complexity, and their interaction. Term significance was obtained from ANOVA, and variance contributions were decomposed by comparing *R*^2^ from nested models. Second, the non-linear “hump” response along each resource axis was quantified using a conservative peak-height metric computed on observed means (peak at interior levels relative to both endpoints), with 95% confidence intervals obtained by bootstrap resampling observations within each curve (i.e., within each fixed level of the other resource).

Compositional variability, computed as described above with vegan as the mean Bray-Curtis distance to the group centroid, was correlated with mean Shannon diversity per cell in the resource grid using Pearson and Spearman correlation tests. The associated scatterplot used an ordinary least-squares fit with 95% confidence bands for visualization (Fig. S7).

Portions of the analysis code were refactored with the assistance of ChatGPT (GPT-5.2 Thinking; OpenAI, accessed February 2026). All code and outputs were verified by the authors.

## Results

### Level of total carbon availability drives community dynamics

Serially transferring our 16-species community for 16 days in 96 resource environments (8 carbon levels × 12 complexity levels) and using an in-house, previously undescribed amplicon library preparation protocol (based on the TaggiMatrix (Adapterama II) method of Glenn *et al.* (2019)^46^, see Supplementary methods, primers are listed in Supplementary Tables S12 & S13), we observed that both resource dimensions significantly influenced community composition; however, variation along the total carbon availability (total-C) gradient explained substantially more compositional variation than along the resource complexity (RC) gradient (PERMANOVA, marginal effects: total-C *F*(1,765) = 968.9, *R*^2^ = 0.546, *p* = 0.0001; RC *F*(1,765) = 39.5, *R*^2^ = 0.022, *p* = 0.0001; Table S3; Fig.02). While numerous lower-abundance taxa exhibited gradual changes in frequency across both gradients (Fig. S2), overall community dynamics were dominated by three species, whose relative abundances shifted sharply along the TCA axis (Fig.02). At the low-total-C end of the gradient (0.03125 g C L⁻¹, 0.0625 g C L⁻¹, 0.0125 g C L⁻¹), *Aeromonas caviae* reached the highest abundance, while at high total-C (1 g C L⁻¹, 2 g C L⁻¹, 4 g C L⁻¹) *Citrobacter koseri* dominated community composition. In the intermediate zone (0.25 g C L⁻¹, 0.5 g C L⁻¹) *Pseudomonas chlororaphis* rises to high abundance, alongside *A. caviae*.

**Figure 02.**
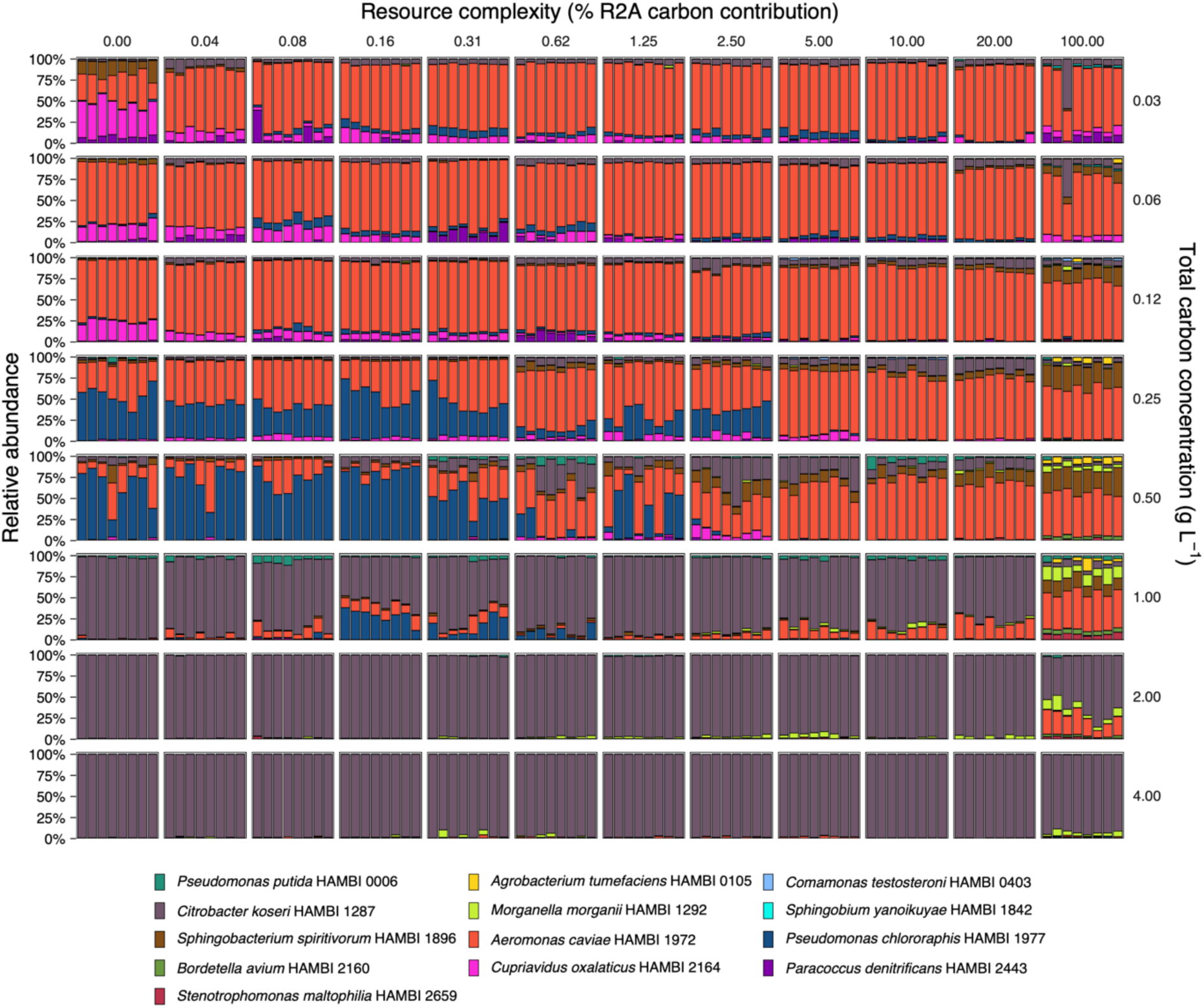
Relative abundance of species in community in all resource environments after 16 day serial passage experiment. Each bar chart panel represents the composition of a serially transferred microbial community in relative abundances after 16 days of passaging in the 96 different resource environments. The total carbon gradient is represented on the *y*-axis, the resource complexity gradient is represented on the *x*-axis. The figure contains 768 data points (8 replicates in 96 resources). Alt text. Bar charts representing the relative abundance of species in the community after the sixteen days of the serial passaging experiment, with one chart for each of the ninety-six resource environments.

**Figure 03.**
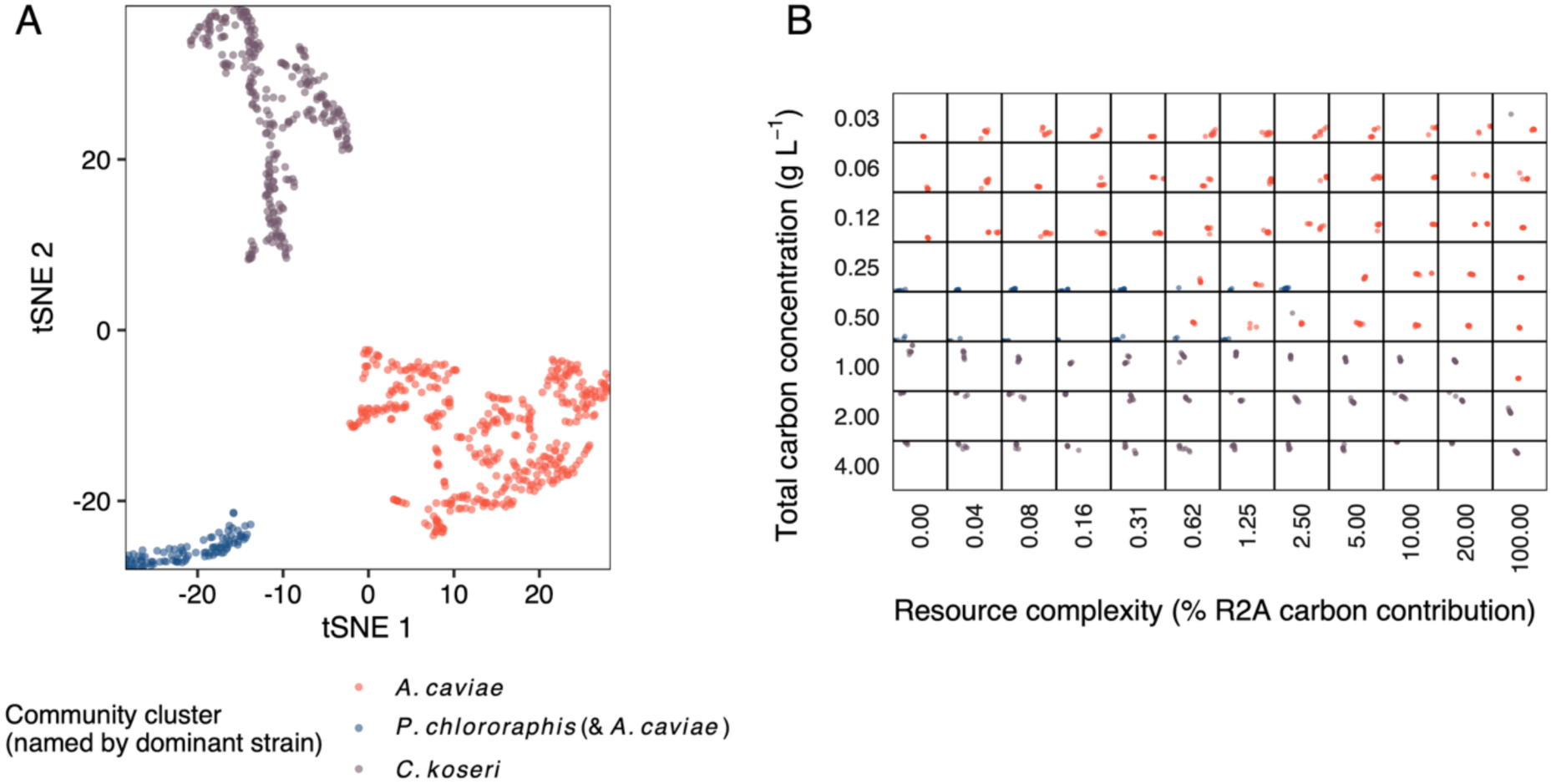
tSNE visualisation (ordination) of 16S rRNA data of community after 16 day serial passage experiment. A | Global clustering of community into three states. These correspond to communities dominated by *A. caviae* (orange), co-dominated by *P. chlororaphis* (dark blue) and *A. caviae*, or dominated by *C. koseri* (grey). The figure contains 768 data points (8 replicates in 96 resources). B | Same ordination partitioned by resource levels. Alt text. Two-panel graphical representation of tSNE ordination analysis, with panel A containing data from all resource environments, and panel B consisting of ninety-six subgraphs, one for each resource environment.

We found that this change in key species along the total-C gradient was not driven by changes in community biomass (Fig. S3), e.g. the low-total-C dominator (*A. caviae*) did not stay at the same absolute abundance along the total-C gradient and only decreased in relative abundance as the high-total-C dominant species (*C. koseri*) took over. Instead, *A. caviae* was lost at both levels.

Ordination analysis revealed that communities clustered into three major compositional states corresponding to dominance by *A. caviae*, co-dominance by *P. chlororaphis* and *A. caviae*, or dominance by *C. koseri* (Fig.03A). These clusters were primarily structured by the total-C gradient, reflecting discrete switches in dominance among the three species. Partitioning the same ordination by resource levels showed that resource complexity modulated these primary dynamics: the co-dominant *P. chlororaphis–A. caviae* state occurred only at low levels of resource complexity, whereas high resource complexity favored exclusive dominance states (Fig.03B). Notably, lowest repeatability between the replicate communities (highest community divergence from group centroid) was observed in the transition zone of the three states (Fig. S4).

In summary, we observed the emergence of a tristable pattern in the serially transferred multispecies synthetic community. The dominant species in the three clusters (*A. caviae* - low total-C, *P. chlororaphis* - intermediate total-C, *C. koseri* - high total-C) are generalists regarding resource metabolism.^43,44^ Together, these results indicate that total carbon availability was the principal driver of community-level state transitions, with resource complexity exerting a secondary, modulatory effect.

### Monoculture lag phase duration explains community outcome

Having observed the partitioning of the community composition into the three clusters along the total-C gradient, we next examined factors that could explain the primary dynamics of the three dominant species (*A. caviae*, *C. koseri*, *P. chlororaphis*; Fig.04A). We cultured the most abundant species of the community experiment at each total-C level but without added complexity, performing three serial transfers to acclimate species to each resource level. In monocultures, these three species differed significantly in lag time in a total-C-dependent manner (Fig.04B; linear model, species×total-C interaction: *F*(2,42) = 9.42, *p* < 0.001), but not in maximum growth rate, carrying capacity, or area under the growth curve (AUC; species×total-C interactions: *F*(2,42) = 1.50, *p* = 0.236; *F*(2,42) = 0.46, *p* = 0.64; and *F*(2,42) = 1.25, *p* = 0.30, respectively; Table S4). Species also differed in maximum growth rate and AUC in a total-C-independent manner (species main effects: *F*(2,44) = 19.66, *p* < 0.001 and *F*(2,44) = 3.43, *p* = 0.041, respectively), but not in lag time or carrying capacity (species main effects: *F*(2,44) = 0.86, *p* = 0.43 and *F*(2,44) = 2.47, *p* = 0.096, respectively). Maximum growth rate ranked consistently across total-C levels as *P. chlororaphis > A. caviae > C. koseri* (Tukey-adjusted pairwise contrasts, all *p* < 0.02; Table S5). AUC showed the same species ordering, but differences were weaker and did not reach significance in pairwise comparisons (Table S6). However, this ranking was inconsistent with community outcomes, where mostly *A. caviae* or *C. koseri* dominated depending strongly on total-C level. These results indicate that lag time was particularly influenced by total-C level in monocultures.

We next tested whether monoculture growth traits predicted community dominance across total-C levels. Across the total-C gradient, shorter lag time was the strongest independent predictor of the abundance of the three dominant species (unique *R*^2^ = 0.314, *p* < 0.001), explaining over twice as much additional variance as higher maximum growth rate (unique *R*^2^ = 0.136, *p* < 0.001), while higher AUC contributed little (unique *R*^2^ *=* 0.039; Table S7). In the intermediate species transition zone (0.25-1.0 g C L⁻¹), lag remained the single strongest independent predictor (unique *R*^2^ = 0.439, *p* < 0.001), but maximum growth rate contributed nearly as much additional variance (unique *R*^2^ = 0.349), indicating that both traits shaped dominance under these conditions (Fig.04C).

When considered alone, lag explained less variation in the transition zone (*R*^2^ = 0.112) than across the full gradient (*R*^2^ = 0.161), consistent with reduced divergence in lag times (Fig.04B). In contrast, the standalone predictive power of growth rate increased in the transition zone (*R*^2^ = 0.083 vs 0.018). Thus, while lag remained central to competitive success, its dominance as the primary predictor weakened as lag times converged and growth rate became comparatively more influential.

Overall, monoculture growth traits had a moderate ability to quantitatively predict the major shifts in community dominance across the total-C gradient, with lag time structuring outcomes at carbon extremes and growth rate becoming comparably important as lag divergence decreased.

### Relationship between resource level and diversity depends on resource complexity

Both gradients (total-C, resource complexity) showed strong, positive, and approximately linear relationships with community biomass (Fig. S5). Biomass was dominated by total-C availability, with smaller but significant contributions from resource complexity and their interaction (linear model: total-C *F*(1,764) = 878.7, *p* < 0.001; RC *F*(1,764) = 51.8, *p* < 0.001; total-C×RC *F*(1,764) = 102.3, *p* < 0.001), together explaining 57.1 % of the variance in biomass (Tables S8 & S9).

In contrast, Shannon diversity responded to the two resource axes in qualitatively different ways (Fig.05). Along the total-C gradient, diversity declined sharply at high concentrations across all levels of resource complexity (Fig.05A), reflecting dominance by *C. koseri*, with a peak at intermediate total-C levels. Along the resource complexity gradient, diversity was comparatively stable across most levels and increased only at the highest complexity level. The non-linear diversity response along the total-C gradient was quantified using a conservative hump metric (Fig.05B), which was positive with 95 % bootstrap confidence intervals excluding zero across all RC levels except 0 %, confirming consistent peak diversity at intermediate total-C concentrations. Thus, total-C primarily imposed a non-linear constraint on diversity, whereas resources complexity buffered diversity across a broad range of levels.

**Figure 04.**
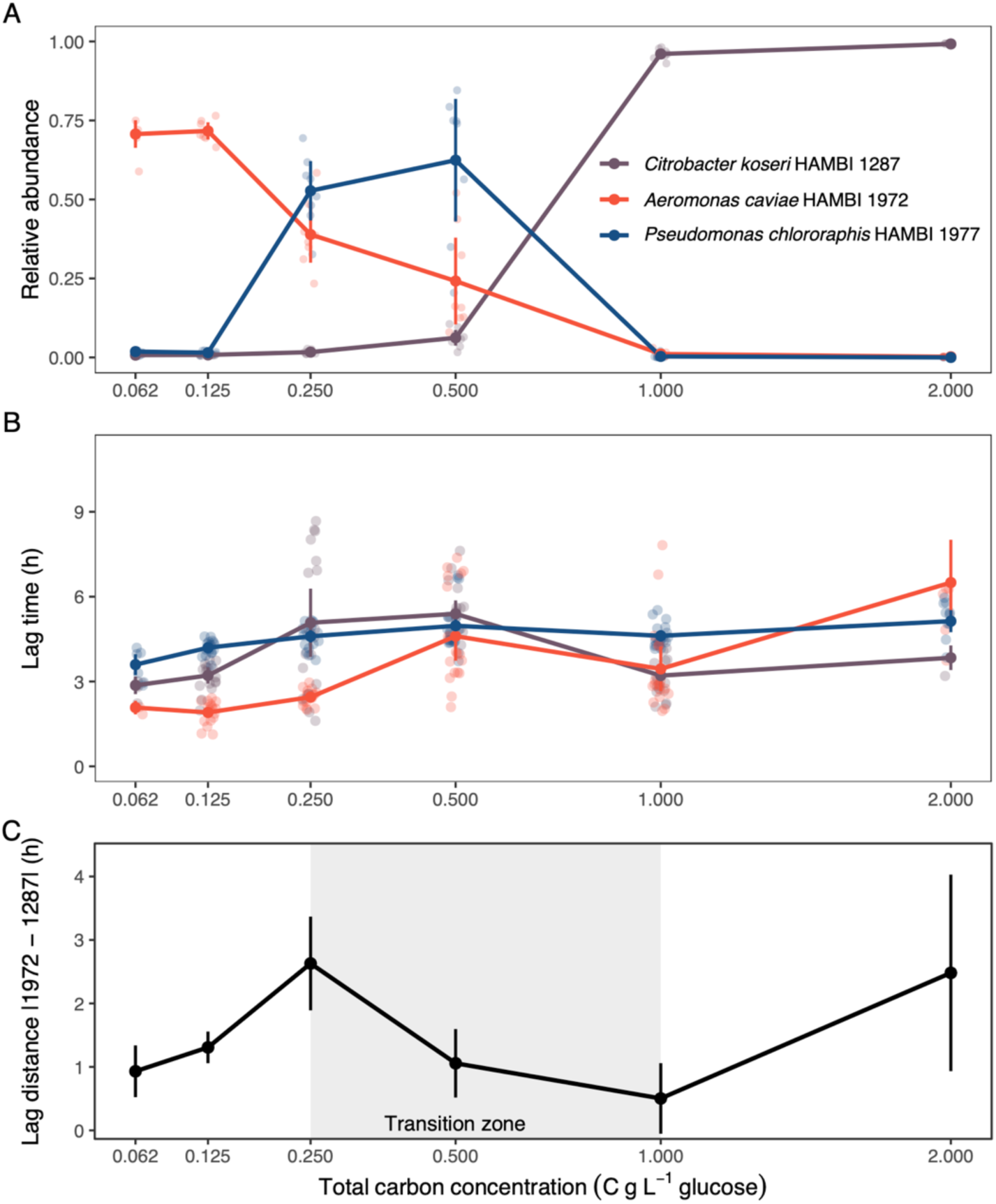
Three-panel figure detailing monoculture growth traits of three dominant species along carbon gradient (mean ± 95 % confidence intervals). A | Relative abundance of species (*N* = 8 replicates per species per time point). B | Duration of lag time of each species at matching carbon levels (*N* = 12–18 per species per time point). C | Distance between lag phase duration between the two most abundant species. Alt text. Three-panel figure detailing monoculture growth traits of the three dominant species along a carbon gradient, with panel A showing the relative abundance of species, panel B showing the duration of lag time of each species in each carbon level, and panel C showing the distance between lag phase duration between the two most abundant species.

**Figure 05.**
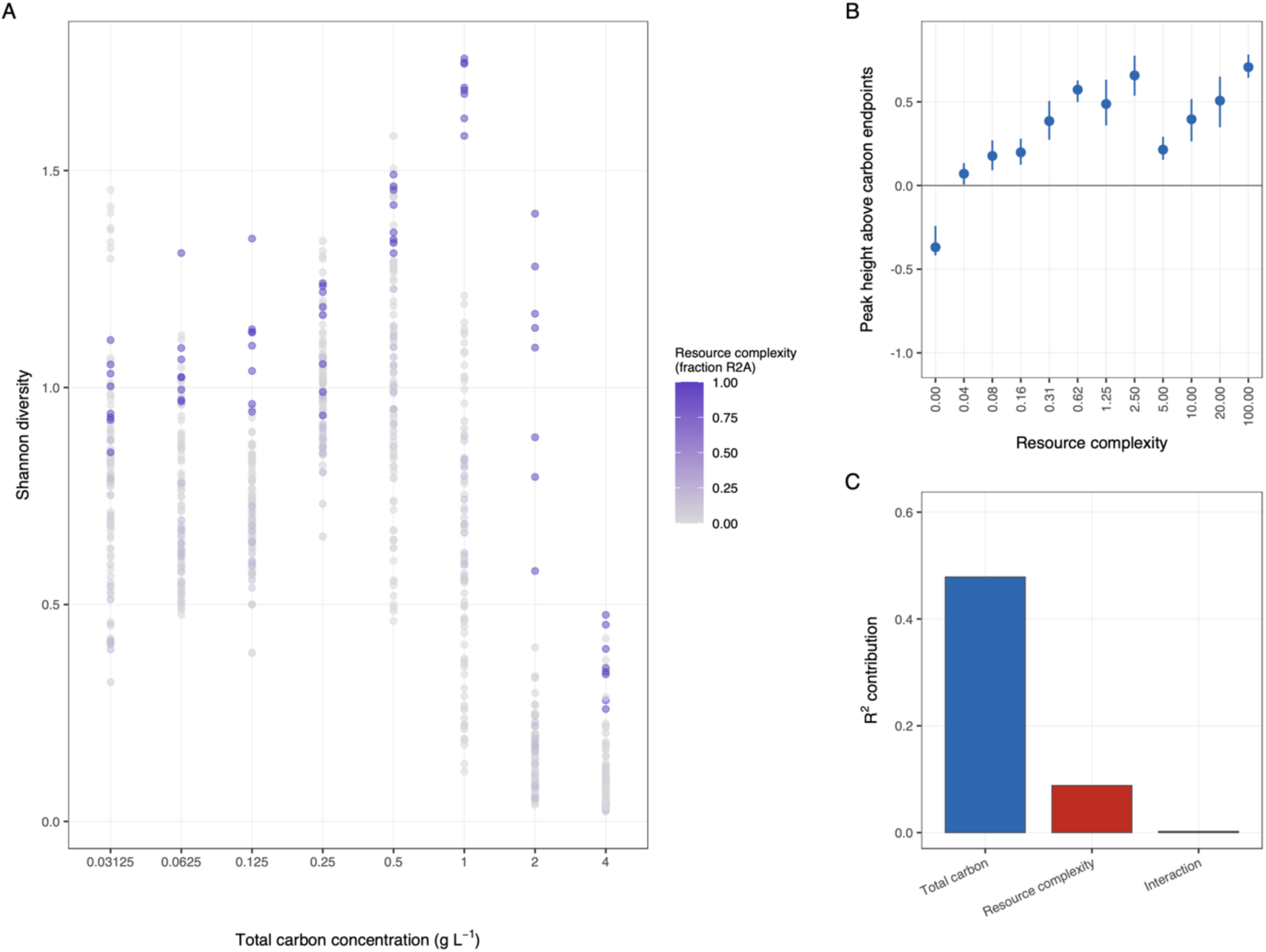
Diversity response along resource gradients. A | Shannon diversity shown separately for each resource environment. The *x-*axis shows the total carbon level (doubling concentrations spaced evenly) and point color indicates resource complexity level. The figure contains 768 data points (8 replicates in 96 resources). B | Hump metric for total carbon gradient (*y*-axis) at each level of resource complexity (*x*-axis). Points show the estimated peak height (hump statistic); error bars show 95 % bootstrap confidence intervals. C | Variance partitioning of Shannon diversity using nested linear models. Bars show unique *R*^2^ contributions of total carbon level, resource complexity (% R2A carbon contribution), and their interaction. Alt text. Three-panel figure representing diversity measures, with panel A containing data on Shannon diversity, resolved for each resource environment, panel showing a hump metric for total carbon gradient (*y*-axis) at each level of resource complexity (*x*-axis), and panel C showing variance partitioning of Shannon diversity using nested linear models.

These patterns were reflected in statistical models: total-C explained the largest share of variation in Shannon diversity (linear *model: F*(1,765) = 846.0*, p* < 0.001; *R*^2^ = 0.479), followed by a smaller but significant effect of resource complexity (*F*(1,765) = 156.0, *p* < 0.001; *R*^2^ = 0.088; Fig.05C; Tables S10 & S11). The total-C×RC interaction was statistically significant but accounted for little additional variance (*F*(1,764) = 4.15, *p* = 0.042; *R*^2^ = 0.002), indicating that the two resources act largely independently.

Furthermore, mean Shannon diversity was strongly correlated with community dispersion across the resource grid (Pearson *r* = 0.67, Spearman *ρ* = 0.82, both *p* < 0.001; Figs. S6 & S7), indicating that conditions supporting higher among-replicate compositional variability (transition zone between dominant species regimes; Figs. 02 & 03) also exhibited greater within-community diversity.

## Discussion

We serially transferred a 16-species synthetic microbial community in a broad range of resource environments for a period of 16 days and observed, consistently over replicates, the emergence of a tristable community state pattern along the total-C gradient. Thus, total-C availability was the principal driver of community-level regime shifts. This is evidenced by the strong, positive linear relationship between total-C and community biomass, and the non-linear impact of total-C on diversity (sharply declining at high concentrations and peaking at intermediate levels). Resource complexity had a smaller but significant effect on community dynamics, particularly in buffering diversity across a broad range of total-C levels. Both resource gradients were shown to act independently in shaping community composition and diversity. The interaction between total-C and resource complexity explained little additional variance, indicating that each gradient had its own unique impact.

Our results underscore the profound influence of total carbon availability on microbial community composition and diversity. Along the total-C gradient, we observed discrete shifts between three dominant community states, accompanied by a strong increase in biomass and a non-linear response of diversity. This combination highlights that resource quantity can simultaneously intensify productivity while constraining diversity, revealing a complex relationship between resource supply and community structure that departs from simple linear expectations.^8^ Glucose, the principal carbon source in the total-C gradient, likely represents a key niche axis for the dominant species (*A. caviae*, *P. chlororaphis*, and *C. koseri*), whereas increasing proportions of R2A-derived carbon appear to buffer diversity across carbon levels without fundamentally restructuring community states.

These findings are consistent with ecological theory predicting that resource gradients can drive community shifts through competitive exclusion or coexistence.^9^ They also align with the framework proposed by Gonze et al. (2017), in which alternative community compositions can arise in response to gradual environmental change. The tristable pattern we observed emerges along a continuous carbon gradient, supporting the notion that shifts in resource supply alone can reorganize community states without changes in the species pool or stochastic colonization history. At the same time, the clustering of communities into discrete compositional types mirrors previous observations that microbial communities often occupy distinct states rather than forming smooth continua.^27,29,53^ Our data suggest that such clustering can arise from the competitive dynamics of a few dominant taxa interacting with resource supply.

The broader literature further contextualizes these patterns. Dal Bello et al. (2021)^16^ showed that increasing resource complexity can increase bacterial richness, though often less than predicted by simple niche-expansion models, while Goldford et al. (2018)^14^ demonstrated that the identity of the supplied carbon source strongly determines community composition at higher taxonomic levels. Together with our results, this suggests that resource quantity structures primary community states, whereas resource complexity modulates diversity within those states. Finally, the fact that dominance shifts could be partially predicted from monoculture growth traits indicates that, even in multispecies settings where cross-feeding and environmental modification can occur,^54^ species-level life-history differences along environmental gradients can retain a measurable imprint on competitive sorting.

Monoculture growth traits provided a mechanistic link between individual performance and community dominance, but their relative importance depended on total carbon availability. Both lag phase and maximal growth rate are known to influence microbial competition,^23^ yet our results show that their predictive value is environmentally contingent. Lag phase, reflecting physiological adjustment to novel environments, can generate strong priority effects in serial-transfer systems, where early starters gain disproportionate access to resources.^24^ Under carbon-poor conditions, such priority effects are likely amplified because early growth rapidly exhausts limited resources, allowing small differences in lag to determine competitive outcomes and potentially alternative community states.

Although the three dominant strains (*A. caviae*, *P. chlororaphis*, and *C. koseri*) are metabolic generalists,^44^ they likely differ in substrate preferences,^55,56^ and align broadly with contrasting ecological strategies: *Aeromonas* species frequently persist in resource-scarce aquatic habitats,^57–61^ consistent with an oligotrophic tendency,^62,63^ whereas *C. koseri*, a close relative of *Escherichia coli*,^64^ reflects a more copiotrophic strategy^62,63^. Such differentiation may maintain pronounced lag divergence at carbon extremes, where early-start advantages dominate. In contrast, at intermediate carbon supply, where lag differences diminish, competitive sorting shifts toward differences in maximum growth rate as part of overall growth performance.^23^ Together, these results indicate that total carbon availability structures community state transitions by modulating which life-history trait governs dominance: lag phase dominates when environmental conditions preserve strong timing differences, whereas maximum growth rate becomes comparatively more important as lag divergence decreases.

Furthermore, our findings highlight the differential effects of resource complexity on diversity and composition. While the total-C gradient primarily imposed a non-linear constraint on diversity, leading to a peak at intermediate concentrations, resource complexity buffered diversity across a broad range of levels. This observation underscores the potential for resource complexity to provide a degree of stability and resilience to microbial communities, particularly in environments where simple resources may be limiting or variable^15,65^. Adopted from plant biology, the stress gradient hypothesis (SGH) proposes that the composition and structure of (microbial) communities are affected by a gradient of environmental stressors by modulating the interactions between the organisms (low stress → facilitation, high stress → competition) ^31,32^. In our data, one species (*C. koseri*) competitively excludes most other species in the high total-C-low diversity cluster consistent with SGH holding that interactions change from facilitative to competitive at increasing resource availability. However, this only holds for the total-C gradient and is not the case with increasing resource complexity resource. This is likely because increasing total-C levels together with increasing resource complexity maintain diversity by ameliorating competition.

In conclusion, our results provide new insights into the complex dynamics of microbial communities under varying resource conditions. By examining the impact of total-C availability and resource complexity on community composition and diversity, we can better appreciate the intricate relationships between resource quantity and community structure. These findings have important implications for our understanding of microbial ecology and for developing more realistic predictive models of ecosystem functioning.

## Data Availability Statement

The 16S rRNA amplicon sequencing data generated in this study have been deposited in the European Nucleotide Archive (ENA) under accession number PRJEB108454. All downstream data tables and code needed to reproduce the analyses will be deposited in Zenodo.

## CRediT authorship contribution statement

AMB – conceptualisation, methodology, project administration, investigation, writing: original draft, writing: editing & reviewing, funding acquisition.

JC – conceptualisation, methodology, project administration, data curation, formal analysis, resources, software, visualisation, investigation, writing: original draft, writing: editing & reviewing.

IKA – investigation, writing: original draft, writing: editing & reviewing.

SP – investigation, writing: original draft.

ML – investigation; writing: original draft.

VM – conceptualisation, visualisation, investigation, writing: editing & reviewing.

TH – conceptualisation, methodology, visualisation, investigation, writing: editing & reviewing, funding acquisition.

## Acknowledgments

AMB thanks Alex Hall for provision of office space and him and his group for discussions around the experiments.

## References

1. Ducklow, H., ‘Microbial Services: Challenges for Microbial Ecologists in a Changing World’, Aquatic Microbial Ecology, 53 (2008), 13–9, 10.3354/ame01220.

2. Orcutt, B. N., et al., ‘Impacts of Deep-sea Mining on Microbial Ecosystem Services’, Limnology and Oceanography, 65/7 (2020), 1489–510, 10.1002/lno.11403.

3. Widder, S., et al., ‘Challenges in Microbial Ecology: Building Predictive Understanding of Community Function and Dynamics’, The ISME Journal, 10/11 (2016), 2557–68, 10.1038/ismej.2016.45.

4. VanEvery, H., et al., ‘Microbiome Epidemiology and Association Studies in Human Health’, Nature Reviews Genetics, 24/2 (2023), 109–24, 10.1038/s41576-022-00529-x.

5. Louw, N. L., et al., ‘Microbiome Assembly in Fermented Foods’, Annual Review of Microbiology, 77/1 (2023), 381–402, 10.1146/annurev-micro-032521-041956.

6. Wagner, M., et al., ‘Microbial Community Composition and Function in Wastewater Treatment Plants’, Antonie van Leeuwenhoek (Netherlands), 81 (2002), 665–80.

7. Verma, S., and A. Kuila, ‘Bioremediation of Heavy Metals by Microbial Process’, Environmental Technology & Innovation, 14 (2019), 100369, 10.1016/j.eti.2019.100369.

8. Hardin, G., ‘The Competitive Exclusion Principle’, Science, New Series, 131/3409 (1960), 1292–7.

9. Tilman, D., Resource Competition and Community Structure. (Princeton, 1982), 10.1515/9780691209654.

10. Ghoul, M., and S. Mitri, ‘The Ecology and Evolution of Microbial Competition’, Trends in Microbiology, 24/10 (2016), 833–45, 10.1016/j.tim.2016.06.011.

11. Estrela, S., et al., ‘Nutrient Dominance Governs the Assembly of Microbial Communities in Mixed Nutrient Environments’, eLife, 10 (2021), e65948, 10.7554/eLife.65948.

12. D’Souza, G., et al., ‘Ecology and Evolution of Metabolic Cross-Feeding Interactions in Bacteria’, Natural Product Reports, 35/5 (2018), 455–88, 10.1039/C8NP00009C.

13. Gralka, M., et al., ‘Trophic Interactions and the Drivers of Microbial Community Assembly’, Current Biology, 30/19 (2020), R1176–88, 10.1016/j.cub.2020.08.007.

14. Goldford, J. E., et al., ‘Emergent Simplicity in Microbial Community Assembly’, Science, 361/6401 (2018), 469–74, 10.1126/science.aat1168.

15. Silverstein, M. R., J. M. Bhatnagar, and D. Segrè, ‘Metabolic Complexity Drives Divergence in Microbial Communities’, Nature Ecology & Evolution, 8/8 (2024), 1493–504, 10.1038/s41559-024-02440-6.

16. Dal Bello, M., et al., ‘Resource–Diversity Relationships in Bacterial Communities Reflect the Network Structure of Microbial Metabolism’, Nature Ecology & Evolution, 5/10 (2021), 1424–34, 10.1038/s41559-021-01535-8.

17. Pacheco, A. R., M. L. Osborne, and D. Segrè, ‘Non-Additive Microbial Community Responses to Environmental Complexity’, Nature Communications, 12/1 (2021), 2365, 10.1038/s41467-021-22426-3.

18. Philippot, L., B. S. Griffiths, and S. Langenheder, ‘Microbial Community Resilience across Ecosystems and Multiple Disturbances’, Microbiology and Molecular Biology Reviews, 85/2 (2021), e00026–20, 10.1128/MMBR.00026-20.

19. Cerulus, B., et al., ‘Transition between Fermentation and Respiration Determines History-Dependent Behavior in Fluctuating Carbon Sources’, eLife, 7 (2018), e39234, 10.7554/eLife.39234.

20. Rolfe, M. D., et al., ‘Lag Phase Is a Distinct Growth Phase That Prepares Bacteria for Exponential Growth and Involves Transient Metal Accumulation’, Journal of Bacteriology, 194/3 (2012), 686–701, 10.1128/JB.06112-11.

21. Bertrand, R. L., ‘Lag Phase Is a Dynamic, Organized, Adaptive, and Evolvable Period That Prepares Bacteria for Cell Division’, Journal of Bacteriology, 201/7 (2019), 10.1128/JB.00697-18.

22. Vermeersch, L., et al., ‘On the Duration of the Microbial Lag Phase’, Current Genetics, 65/3 (2019), 721–7, 10.1007/s00294-019-00938-2.

23. Aranda-Díaz, A., et al., ‘Assembly of Stool-Derived Bacterial Communities Follows “Early-Bird” Resource Utilization Dynamics’, Cell Systems, 16/4 (2025), 101240, 10.1016/j.cels.2025.101240.

24. Chu, D., and D. J. Barnes, ‘The Lag-Phase during Diauxic Growth Is a Trade-off between Fast Adaptation and High Growth Rate’, Scientific Reports, 6/1 (2016), 25191, 10.1038/srep25191.

25. Manhart, M., and E. I. Shakhnovich, ‘Growth Tradeoffs Produce Complex Microbial Communities on a Single Limiting Resource’, Nature Communications, 9/1 (2018), 3214, 10.1038/s41467-018-05703-6.

26. Lenski, R. E., and M. Travisano, ‘Dynamics of Adaptation and Diversification: A 10,000-Generation Experiment with Bacterial Populations.’, Proceedings of the National Academy of Sciences, 91/15 (1994), 6808–14, 10.1073/pnas.91.15.6808.

27. Dubinkina, V., et al., ‘Multistability and Regime Shifts in Microbial Communities Explained by Competition for Essential Nutrients’, eLife, 8 (2019), e49720, 10.7554/eLife.49720.

28. Van Dijk, B., et al., ‘Trusting the Hand That Feeds: Microbes Evolve to Anticipate a Serial Transfer Protocol as Individuals or Collectives’, BMC Evolutionary Biology, 19/1 (2019), 201, 10.1186/s12862-019-1512-2.

29. Li, Y., et al., ‘Moisture-Driven Microbial Regime Shifts Mediate Nutrient Dynamics in Reservoir Riparian Zones’, Water Research, 287 (2025), 124309, 10.1016/j.watres.2025.124309.

30. Wright, E. S., and Vetsigian, Kalin H, ‘Inhibitory Interactions Promote Frequent Bistability among Competing Bacteria’, Nature Communications, 7/11274 (2016), ncomms11274, 10.1038/ncomms11274.

31. Bertness, M. D., and R. Callaway, ‘Positive Interactions in Communities’, Trends in Ecology & Evolution, 9/5 (1994), 191–3, 10.1016/0169-5347(94)90088-4.

32. Fraser, L. H., et al., ‘Worldwide Evidence of a Unimodal Relationship between Productivity and Plant Species Richness’, Science, 349/6245 (2015), 302–5, 10.1126/science.aab3916.

33. Piccardi, P., B. Vessman, and S. Mitri, ‘Toxicity Drives Facilitation between 4 Bacterial Species’, Proceedings of the National Academy of Sciences, 116/32 (2019), 15979–84, 10.1073/pnas.1906172116.

34. Harris, K. J., and A. E. Bennett, ‘Exploring Bacterial Interactions Under the Stress Gradient Hypothesis in Response to Selenium Stress’, Environmental Microbiology Reports, 17/5 (2025), e70191, 10.1111/1758-2229.70191.

35. Hernandez, D. J., et al., ‘Environmental Stress Destabilizes Microbial Networks’, The ISME Journal, 15/6 (2021), 1722–34, 10.1038/s41396-020-00882-x.

36. Lamont, B. B., ‘The Species Richness–Resource Availability Relationship Is Hump-Shaped’, Perspectives in Plant Ecology, Evolution and Systematics, 65 (2024), 125824, 10.1016/j.ppees.2024.125824.

37. Li, R., et al., ‘Distinct Soil Bacterial Patterns along Narrow and Broad Elevational Gradients in the Grassland of Mt. Tianshan, China’, Scientific Reports, 12/1 (2022), 136, 10.1038/s41598-021-03937-x.

38. Navas-Molina, J. A., et al., ‘The Microbiome and Big Data’, Current Opinion in Systems Biology, 4 (2017), 92–6, 10.1016/j.coisb.2017.07.003.

39. Ringel, Y., et al., ‘High Throughput Sequencing Reveals Distinct Microbial Populations within the Mucosal and Luminal Niches in Healthy Individuals’, Gut Microbes, 6/3 (2015), 173–81, 10.1080/19490976.2015.1044711.

40. Di Bella, J. M., et al., ‘High Throughput Sequencing Methods and Analysis for Microbiome Research’, Journal of Microbiological Methods, 95/3 (2013), 401–14, 10.1016/j.mimet.2013.08.011.

41. Diaz-Colunga, J., et al., ‘Top-down and Bottom-up Cohesiveness in Microbial Community Coalescence’, Proceedings of the National Academy of Sciences, 119/6 (2022), e2111261119, 10.1073/pnas.2111261119.

42. Goldford, J. E., et al., ‘Emergent Simplicity in Microbial Community Assembly’, Science, 361/6401 (2018), 469–74, 10.1126/science.aat1168.

43. Cairns, J., et al., ‘Pre-Exposure of Abundant Species to Disturbance Improves Resilience in Microbial Metacommunities’, Nature Ecology & Evolution, 9/3 (2025), 395–405, 10.1038/s41559-024-02624-0.

44. Cairns, J., et al., ‘Construction and Characterization of Synthetic Bacterial Community for Experimental Ecology and Evolution’, Frontiers in Genetics, 9 (2018), 1–12, 10.3389/fgene.2018.00312.

45. Hogle, S. L., et al., ‘Effects of Phenotypic Variation on Consumer Coexistence and Prey Community Structure’, Ecology Letters (22 Nov. 2021), ele.13924, 10.1111/ele.13924.

46. Glenn, T. C., et al., ‘Adapterama II: Universal Amplicon Sequencing on Illumina Platforms (TaggiMatrix)’, PeerJ, 7 (2019), e7786, 10.7717/peerj.7786.

47. R Core Team, R: A Language and Environment for Statistical Computing. (R Foundation for Statistical Computing, 2024), https://www.R-project.org/.

48. Oksanen J, Simpson G, Blanchet F, Kindt R, Legendre P, Minchin P, et al., Vegan: Community Ecology Package, version R package version 2.6-10 (2025), https://CRAN.R-project.org/package=vegan.

49. Anderson, M. J., ‘A New Method for Non-Parametric Multivariate Analysis of Variance: NON-PARAMETRIC MANOVA FOR ECOLOGY’, Austral Ecology, 26/1 (2001), 32–46, 10.1111/j.1442-9993.2001.01070.pp.x.

50. Maaten, L. van der, and G. Hinton, ‘Visualizing Data Using T-SNE’, Journal of Machine Learning, 9 (2008), 2579–605.

51. Lenth, R., and J. Piaskowski, Estimated Marginal Means, Aka Least-Squares Means., version R package version 2.0.1 (n.d.), https://rvlenth.github.io/emmeans/.

52. Gonze, D., et al., ‘Multi-Stability and the Origin of Microbial Community Types’, The ISME Journal, 11 (2017), 2159–66, 10.1038/ismej.2017.60.

53. Pascual-García, A., et al., ‘Replicating Community Dynamics Reveals How Initial Composition Shapes the Functional Outcomes of Bacterial Communities’, Nature Communications, 16/1 (2025), 3002, 10.1038/s41467-025-57591-2.

54. Santamaria, G., et al., ‘Evolution and Regulation of Microbial Secondary Metabolism’, eLife, 11 (2022), e76119, 10.7554/eLife.76119.

55. Estrela, S., et al., ‘Functional Attractors in Microbial Community Assembly’, Cell Systems, 13/1 (2022), 29–42.e7, 10.1016/j.cels.2021.09.011.

56. Vila, J. C. C., et al., ‘Metabolic Similarity and the Predictability of Microbial Community Assembly’, preprint, 28 Oct. 2023, 10.1101/2023.10.25.564019.

57. Chaix, G., et al., ‘Distinct Aeromonas Populations in Water Column and Associated with Copepods from Estuarine Environment (Seine, France)’, Frontiers in Microbiology, 8 (2017), 1259, 10.3389/fmicb.2017.01259.

58. Simmons, G., et al., ‘Contamination of Potable Roof-Collected Rainwater in Auckland, New Zealand’, Water Research, 35/6 (2001), 1518–24, 10.1016/S0043-1354(00)00420-6.

59. Egorov, A. I., et al., ‘Occurrence of Aeromonas Spp. in a Random Sample of Drinking Water Distribution Systems in the USA’, Journal of Water and Health, 9/4 (2011), 785–98, 10.2166/wh.2011.169.

60. Borchardt, M. A., M. E. Stemper, and J. H. Standridge, ‘*Aeromonas* Isolates from Human Diarrheic Stool and Groundwater Compared by Pulsed-Field Gel Electrophoresis’, Emerging Infectious Diseases, 9/2 (2003), 224–8, 10.3201/eid0902.020031.

61. Razzolini, M. T. P., et al., ‘Aeromonas Presence in Drinking Water from Collective Reservoirs and Wells in Peri-Urban Area in Brazil’, Brazilian Journal of Microbiology, 41/3 (2010), 694–9, 10.1590/S1517-83822010000300020.

62. Soler-Bistué, A., L. L. Couso, and I. E. Sánchez, ‘The Evolving Copiotrophic/Oligotrophic Dichotomy: From Winogradsky to Physiology and Genomics’, Environmental Microbiology, 25/7 (2023), 1232–7, 10.1111/1462-2920.16360.

63. Stone, B. W. G., et al., ‘Life History Strategies among Soil Bacteria—Dichotomy for Few, Continuum for Many’, The ISME Journal, 17/4 (2023), 611–9, 10.1038/s41396-022-01354-0.

64. Jabeen, I., et al., ‘A Brief Insight into Citrobacter Species - a Growing Threat to Public Health’, Frontiers in Antibiotics, 2 (2023), 1276982, 10.3389/frabi.2023.1276982.

65. Smith, R. S., E. L. Johnston, and G. F. Clark, ‘The Role of Habitat Complexity in Community Development Is Mediated by Resource Availability’, PLoS ONE, 9/7 (2014), e102920, 10.1371/journal.pone.0102920.

