## Supplementary Information for "Ecological tristability driven by total carbon availability over resource complexity in a synthetic microbial community"

This file includes

- Supplementary Figures S1-S7
- Supplementary Tables S1-S13
- Supplementary Methods

### Supplementary Methods

#### Supplementary DNA library preparation methods

#### Overview

The in-house library preparation method for constructing 16S amplicon libraries for community composition analysis has been adapted from Adapterama (TaggiMatrix) system[1]. In short, this system uses a two-stage fusion PCR approach with tailed primers to create quadruple-tagged Illumina TruSeq-style amplicon libraries. In the first amplification step (PCR-1), the target is amplified, and each sample is barcoded with a unique combination of forward and reverse tags (combinatorial dual barcoding). The reactions are subsequentially purified and combined into pools of maximum 96 samples. In the second amplification (PCR-2), Illumina compatible i5 and i7 index sequences are added on each pool (unique dual indexing) together with Illumina adapters. The PCR-2 products are purified and size-selected using Carboxyl-modified Sera-Mag Magnetic Speed-beads[2], quantified by a fluorescence-based method (Qubit 2.0, ThermoFisher) and pooled into one library in equal amounts (by DNA mass).

#### DNA extraction

Genomic DNA from the bacterial cells was released by heat lysis. In short, 3 ul bacterial culture was added to 200 ul ultrapure water (MQ-H2O) and mixed well by vortexing. 50-100 µL diluted culture was transferred to a PCR plate, incubated 10 minutes at 100º C in a PCR machine and cooled down by placing on ice. Plate was vortexed for 15 seconds and spun down for 15 seconds. Crude DNA was directly used for PCR) amplification by pipetting from the top to avoid debris.

#### Community control

A control sample with known community composition was added to each batch of DNA extraction (95 samples + 1 community control). Community control sample (CC) was prepared in a large quantity by cultivating each 23 HAMBI species axenically in 100 % R2A at 25° C for 48 or 96 hours. The cultures were concentrated to roughly equal densities, combined in equal volumes, aliquoted and stored at -20° C. The cell density of each axenic culture was simultaneously verified by CFU (colony forming unit) count.

**Primers used**

Locus specific primers for amplifying the V3 region of the 16S rRNA gene of the 23 HAMBI species were designed by Shane Hogle, University of Turku. Sequences of the primers are 16S_HAMBI_341F: 5' – CCTACGGGAGGCAGCAG -3', 16S_HAMBI_518R: 5' – ATTACCGCGGCTGCTGG - 3'. In all other aspects the oligos used are as presented in Glenn *et al* (2019)[1]. To minimize cross contamination happening during oligo synthesis/HPLC purification we chose to use NGS grade oligos (NGSO-Bronze, Sigma Aldrich). For full primer sequences see Supplementary Tables S12 & S13.

**Library preparation protocols (steps 1-9)**

**Step 1. PCR-1 protocol**

| Reagent | Final [c] | [V] per 1 rxn |
|---|---|---|
| 2x My Taq HS Red-mix (Meridian Bioscience/Bioline) | 1x | 10.0 µL |
| Forward primer (fusion primers 1-12 @ 2.5 uM) | 0.3 µM | 2.4 µL |
| Reverse primer (fusion primers A-H @ 2.5 uM) | 0.3 µM | 2.4 µL |
| H2O | | 1.2 µL |
| DNA | | 4.0 µL |
| | Total | 20 µL |

Samples were amplified using Bio-Rad S1000/C1000 and Applied Biosystems 2720 thermocyclers with the following parameters, 95 °C for 1 min, followed by 25 cycles at 95 °C for 15 sec, 58 °C for 15 sec, and 72 °C for 10 sec; and final extension at 72 °C for 5 min, hold at 4 °C. All work prior to PCR amplification was conducted in a designated pre-PCR laboratory and all steps with PCR amplified material were done in a separate post-PCR laboratory space.

**Steps 2-4. Amplification check, purification and pooling.** Proper amplification was checked by quantifying the PCR product by a fluorescence-based method (dsDNA High Sensitivity kit on Qubit 2.0 instrument, Thermo Fisher Scientific). The reactions (15 µL) were immediately purified using Just-a-Plate 96 PCR Normalization and Purification Kit (Charm Biotech) following manufacturer's standard protocol and stored in -20° C. Purified PCR products were pooled in equal volumes, 5-10 µL/reaction.

**Step 5. PCR-2 protocol**

| Reagent | Final [c] | [V] per 1 rxn |
| --- | --- | --- |
| 2x My Taq HS Red-mix (Meridian bioscience/Bioline) | 1x | 10.0 µL |
| Forward primer (fusion primers 1-12 @ 5 µM) | 0.5 µM | 2.0 µL |
| Reverse primer (fusion primers A-H @ 5 µM) | 0.5 µM | 2.0 µL |
| H2O | | 4.0 µL |
| Purified and pooled PCR-1 product | | 2.0 µL |
| | Total | 20 µL |

Samples were amplified using Bio-Rad S1000/C1000 and Applied Biosystems 2720 thermocyclers with the following parameters, 95 °C for 1 min, followed by 10 cycles at 95 °C for 15 sec, 58 °C for 15 sec, and 72 °C for 10 sec; and final extension at 72 °C for 5 min, hold at 4 °C.

**Step 6. SPRI bead clean-up and size selection.** PCR-2 products were cleaned and size selected for c. > 100 bp immediately following amplification, using Carboxyl-modified Sera-Mag Magnetic Speed-beads (Cytiva) in a PEG/NaCl buffer (Rohland & Reich 2012)[2]. A ratio of 1:1 Serapure mixture to PCR-product was used.

**Protocol for SPRI bead clean-up and size selection with Serapure mixture**

Instructions for preparing Serapure mixture, see https://ethanomics.files.wordpress.com/2012/08/serapure_v2-2.pdf.

Mix/vortex Serapure mixture thoroughly (e.g., a Falcon tube sideways on a Vortex Genie 2 with a plate adapter, speed 5-6 for 5 minutes) right before use. General tips: Use molecular grade water for eluation and ethanol dilution. Prepare fresh 80 % ethanol daily. For all steps use filter tips, except autoclaved non-filter tips for ethanol wash steps.

1. Mix equal volumes of Serapure mixture and PCR product (add $H_2O$ to PCR product if needed).
2. Incubate at RT for 5 min (vortex during incubation). Spin down very briefly if needed.
3. Place on magnet until liquid is clear.
4. Remove and discard supernatant (DNA is now attached to beads).
5. Keep on magnet, add 400 µL EtOH (80 %).
6. Incubate > 30 sec.
7. Remove and discard ethanol. Repeat wash (steps 5-7).

8. Allow beads to dry (EtOH evaporates usually in 3-5 min, but do not overdry seen as ”cracking” of the bead pellet). If needed, use 10 µL tips to pipette out residual EtOH.
9. Resuspend in desired volume of water.
10. Vortex until beads are mixed with water.
11. Place on magnet and take out desired volume of supernatant. Make sure no beads are in the supernatant.

**Steps 7-9. DNA quantification, second pooling and final library check**

DNA concentrations of the purified PCR-2 products were measured by a fluorescence-based method (Qubit dsDNA High Sensitivity kit on Qubit 2.0 instrument, Thermo Fisher Scientific). The pools were combined into two final libraries in equal amounts (by DNA mass). The correct size of the final libraries was checked on a Bioanalyzer High Sensitivity DNA kit run (Agilent). The expected size should be c. 320-380 bp and no other (short) fragments should be visible.

The final two libraries were sequenced by Finnish Functional Genomics Centre (Turku Bioscience Centre, Finland) on an Illumina Miseq instrument, on two runs, using 2 x 150 bp read length and MiSeq Reagent Kit v2. Libraries were spiked with 10 or 20 % PhiX Control v3 Library.

### Supplementary Figures

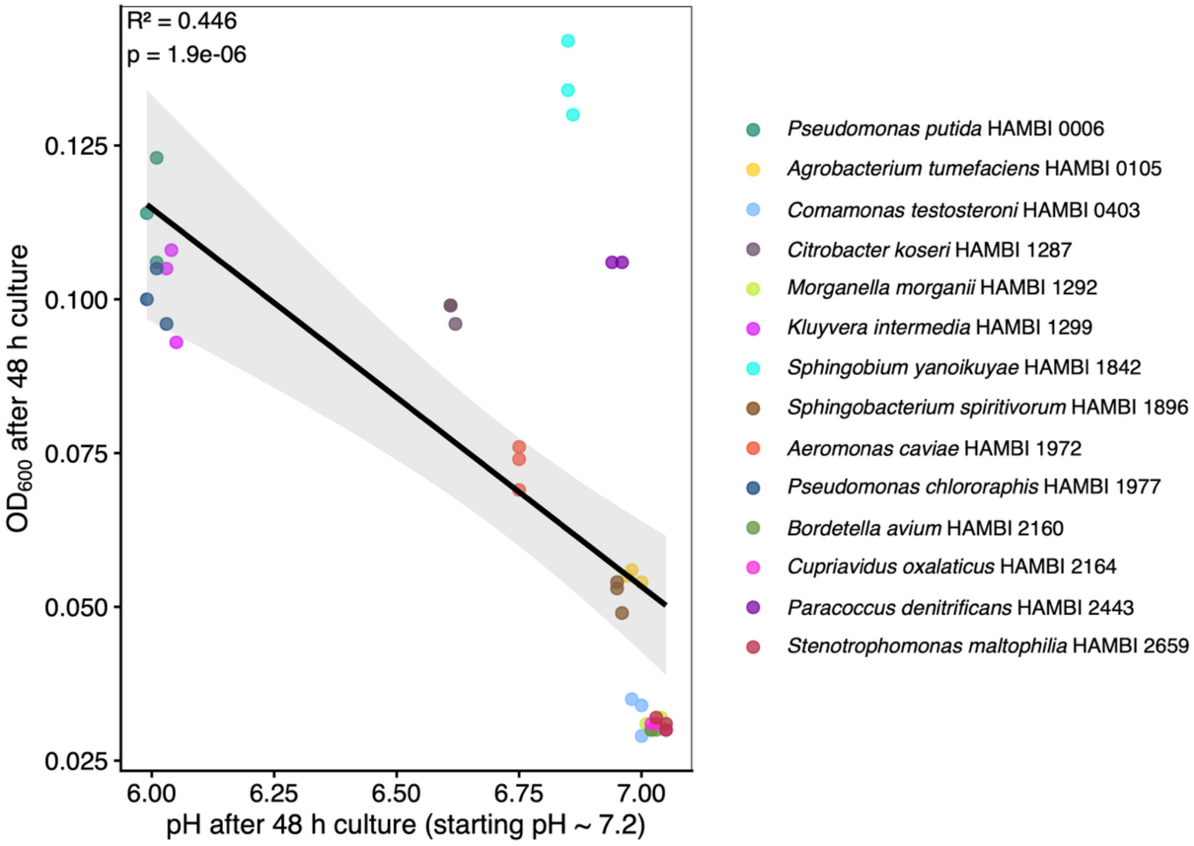


**Supplementary Figure 01. pH-changing potential of 16 species in monoculture at 4 g C $L^{-1}$.** Individual species of the community were assessed for their ability to change pH in monoculture (M9 minimal medium supplemented with 4 g C $L^{-1}$). Each species was tested in triplicate.

Alt text. Dot-and-line graph detailing to which extent each of the sixteen species in the community is able to change the pH of the medium.

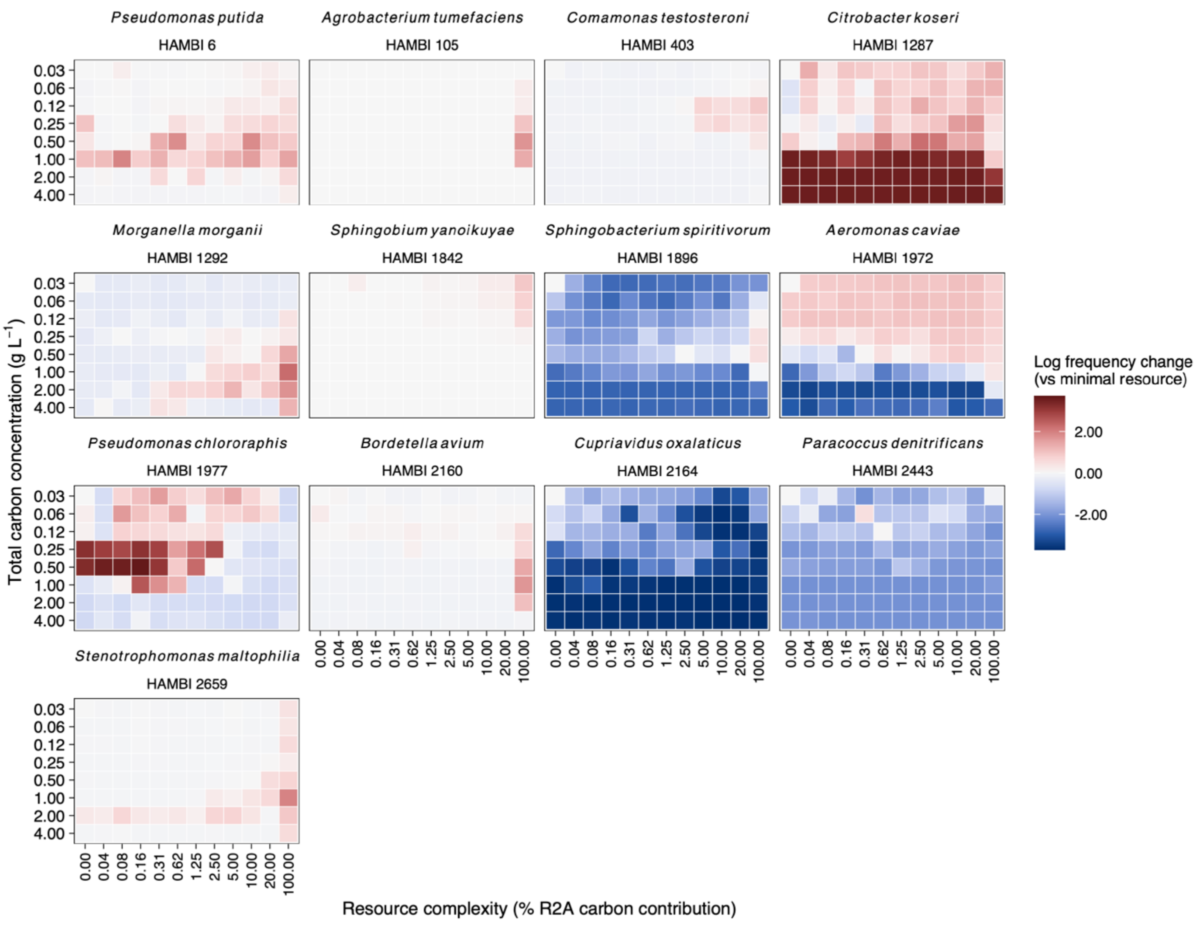


**Supplementary Figure 02. Frequency change of individual species after 16-day serial passage experiment compared to minimal resource conditions.** For each species, the heatmap figure represents changes in relative abundance of the species in each resource environment compared to minimal resource conditions. Red colours indicate increases relative to minimal resource conditions, blue colours indicate decreses compared to minimal resource conditions. The total-C gradient is represented on the *y*-axis, resource complexity is represented on the *x*-axis. The figure contains 768 data points (8 replicates in 96 resources).

Alt text. Heatmap figures for each species representing changes in relative abundance of the species in each resource environment compared to minimal resource conditions.

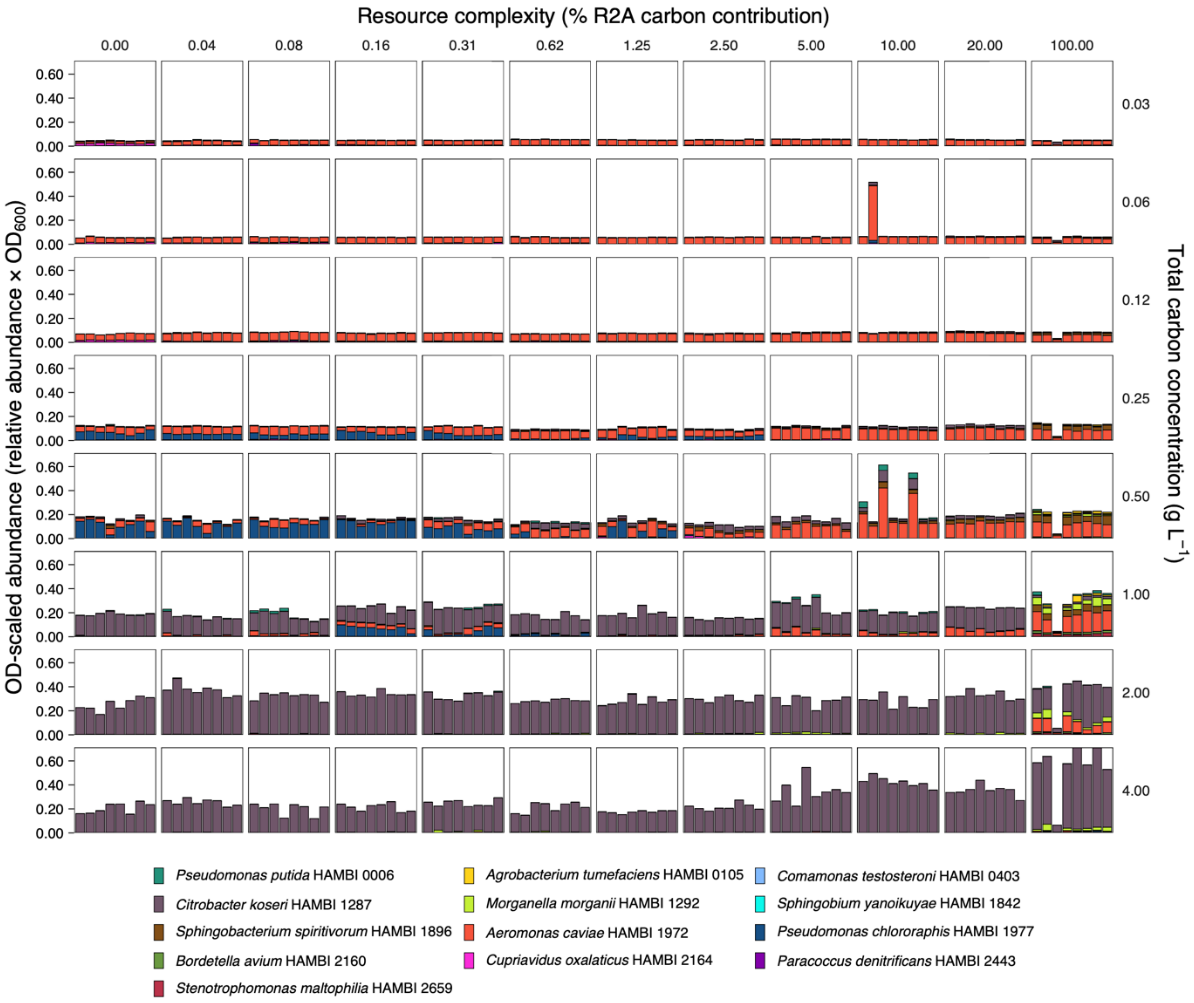


**Supplementary Figure 03. Absolute abundance of species in community in all resource environments after 16-day serial passage experiment.** Each bar chart panel represents the composition of a serially transferred microbial community in absolute abundances after 16 days of passaging. The total-C gradient is represented on the *y*-axis, the resource complexity gradient is represented on the *x*-axis. The figure contains 768 data points (8 replicates in 96 resources).

Alt text. Bar charts representing the absolute abundance of species in the community after the sixteen days of the serial passaging experiment. There is one chart for each of the ninety-six resource environments.

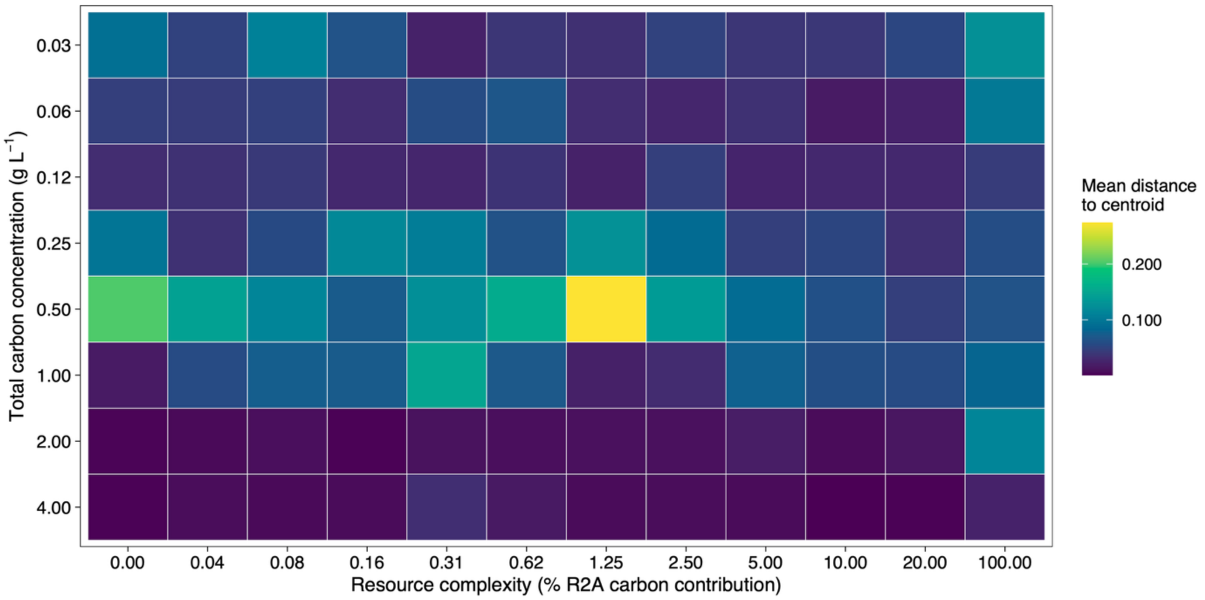


**Supplementary Figure 04. Repeatability between replicate communities is lowest in transition zone of resource gradients.** Heatmap figure detailing the repeatability between the eight replicate communities for each resource environment. Dark blue hues correspond to high repeatability, green and yellow hues correspond to low repeatability. The figure contains 768 data points (8 replicates in 96 resources).

Alt text. Heatmap figure detailing the repeatability between the eight replicate communities for each resource environment.

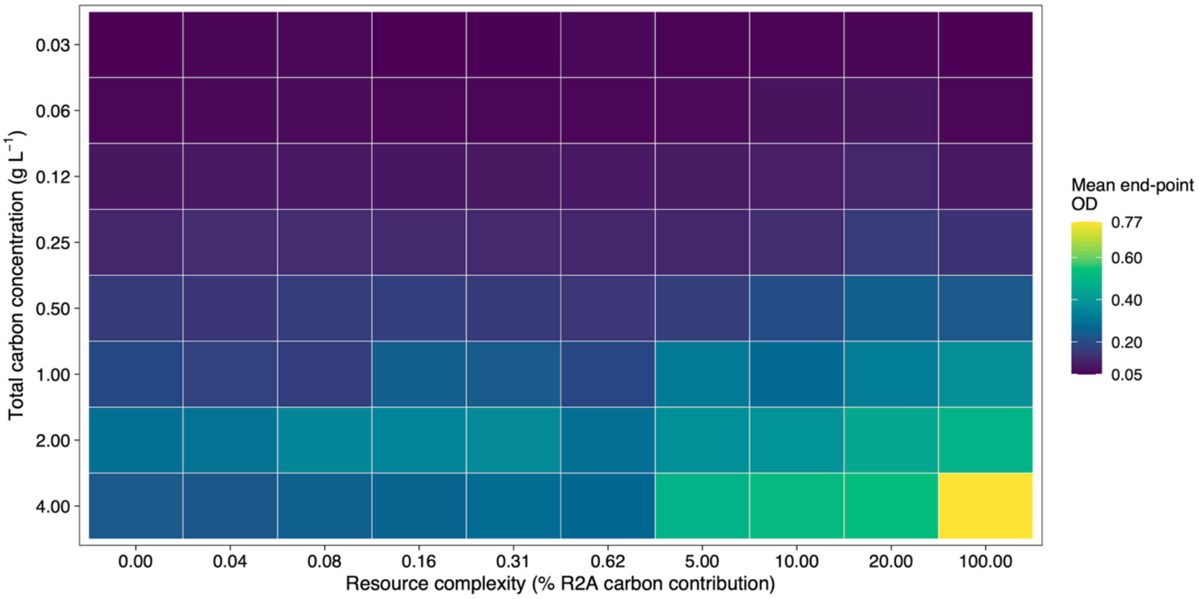


**Supplementary Figure 05. Community biomass in all resource environments after 16-day serial passage experiment.** Heatmap figure detailing the mean end-point biomass of the eight replicate communities in each resource environment. Dark blue hues correspond to low $OD_{600}$, green and yellow hues correspond to high $OD_{600}$**.** The figure contains 768 data points (8 replicates in 96 resources).

Alt text. Heatmap figure detailing the mean end-point biomass of the eight replicate communities in each resource environment.

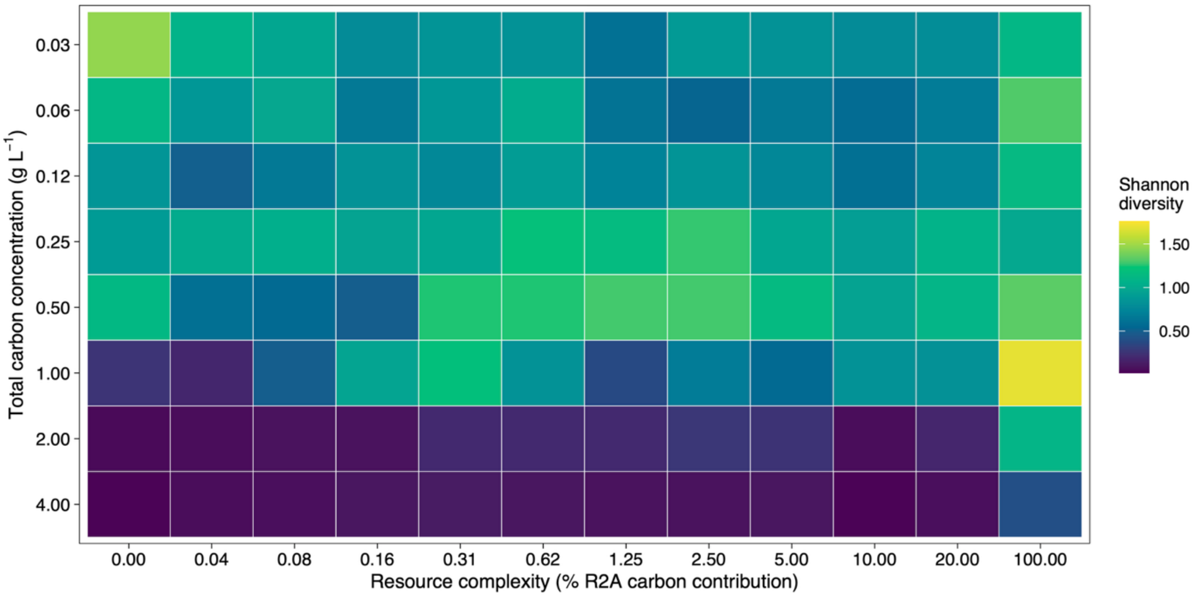


**Supplementary Figure 06. Heatmap of community diversity after 16-day serial passage experiment.** Heatmap figure detailing the mean diversity of the eight replicate communities for each resource environment at the end of the experiment. Dark blue hues correspond to low Shannon diversity, green and yellow hues correspond to high Shannon diversity**.** The figure contains 768 data points (8 replicates in 96 resources).

Alt text. Heatmap figure detailing the mean diversity of the eight replicate communities for each resource environment at the end of the experiment.

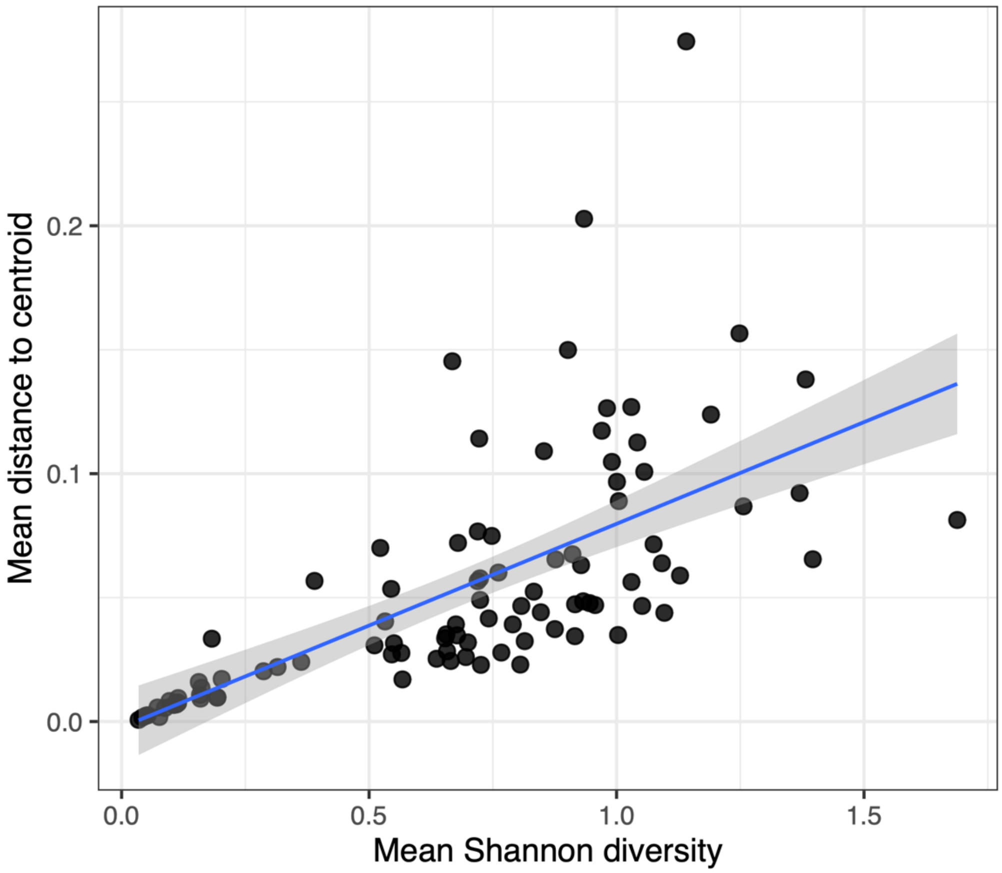


**Supplementary Figure 07. Relationship between diversity and community dispersion.** The figure shows a linear regression line with S.E.M. error bands for 768 data points (8 replicates in 96 resources).

Alt text. Dot-and-line graph visualising the relationship (linear regression) between diversity and community dispersion.

**Supplementary Tables**

**Supplementary Table S1. Carbon composition of experimental media.** Values show the carbon content of all 96 media used in the experiment. Glucose C (g C $L^{-1}$) indicates the carbon added as glucose to M9 minimal medium. R2A carbon (%) refers to the proportion of total carbon derived from R2A powder, including any glucose present in R2A. R2A-derived C (g C $L^{-1}$) indicates the carbon contributed by R2A, and total C (g C $L^{-1}$) is the sum of glucose- and R2A-derived carbon

| **Glucose (g C $L^{-1}$)** | **R2A carbon (%)** | **R2A-derived carbon (g C $L^{-1}$)** | **Total carbon (g C $L^{-1}$)** |
|---|---|---|---|
| 4 | 0 | 0 | 4 |
| 2 | 0 | 0 | 2 |
| 1 | 0 | 0 | 1 |
| 0.50 | 0 | 0 | 0.50 |
| 0.25 | 0 | 0 | 0.25 |
| 0.125 | 0 | 0 | 0.125 |
| 0.062 | 0 | 0 | 0.062 |
| 0.031 | 0 | 0 | 0.031 |
| 3.998 | 0.039 | 0.002 | 4 |
| 1.999 | 0.039 | 0.001 | 2 |
| 1.000 | 0.039 | 0.000 | 1 |
| 0.500 | 0.039 | 0.000 | 0.50 |
| 0.250 | 0.039 | 0.000 | 0.25 |
| 0.125 | 0.039 | 0.000 | 0.125 |
| 0.062 | 0.039 | 0.000 | 0.062 |
| 0.031 | 0.039 | 0.000 | 0.031 |
| 3.997 | 0.078 | 0.003 | 4 |
| 1.998 | 0.078 | 0.002 | 2 |
| 0.999 | 0.078 | 0.001 | 1 |
| 0.500 | 0.078 | 0.000 | 0.50 |
| 0.250 | 0.078 | 0.000 | 0.25 |
| 0.125 | 0.078 | 0.000 | 0.125 |
| 0.062 | 0.078 | 0.000 | 0.062 |
| 0.031 | 0.078 | 0.000 | 0.031 |
| 3.994 | 0.156 | 0.006 | 4 |
| 1.997 | 0.156 | 0.003 | 2 |
| 0.998 | 0.156 | 0.002 | 1 |
| 0.499 | 0.156 | 0.001 | 0.50 |
| 0.250 | 0.156 | 0.000 | 0.25 |

| | | | |
|---|---|---|---|
| 0.125 | 0.156 | 0.000 | 0.125 |
| 0.062 | 0.156 | 0.000 | 0.062 |
| 0.031 | 0.156 | 0.000 | 0.031 |
| 3.987 | 0.312 | 0.012 | 4 |
| 1.994 | 0.312 | 0.006 | 2 |
| 0.997 | 0.312 | 0.003 | 1 |
| 0.498 | 0.312 | 0.002 | 0.50 |
| 0.249 | 0.312 | 0.001 | 0.25 |
| 0.125 | 0.312 | 0.000 | 0.125 |
| 0.062 | 0.312 | 0.000 | 0.062 |
| 0.031 | 0.312 | 0.000 | 0.031 |
| 3.975 | 0.625 | 0.025 | 4 |
| 1.988 | 0.625 | 0.012 | 2 |
| 0.994 | 0.625 | 0.006 | 1 |
| 0.497 | 0.625 | 0.003 | 0.50 |
| 0.248 | 0.625 | 0.002 | 0.25 |
| 0.124 | 0.625 | 0.001 | 0.125 |
| 0.062 | 0.625 | 0.000 | 0.062 |
| 0.031 | 0.625 | 0.000 | 0.031 |
| 3.95 | 1.25 | 0.05 | 4 |
| 1.975 | 1.25 | 0.025 | 2 |
| 0.988 | 1.25 | 0.012 | 1 |
| 0.494 | 1.25 | 0.006 | 0.50 |
| 0.247 | 1.25 | 0.003 | 0.25 |
| 0.123 | 1.25 | 0.002 | 0.125 |
| 0.062 | 1.25 | 0.001 | 0.062 |
| 0.031 | 1.25 | 0.000 | 0.031 |
| 3.90 | 2.50 | 0.10 | 4 |
| 1.95 | 2.50 | 0.05 | 2 |
| 0.975 | 2.50 | 0.025 | 1 |
| 0.487 | 2.50 | 0.012 | 0.50 |
| 0.244 | 2.50 | 0.006 | 0.25 |
| 0.122 | 2.50 | 0.003 | 0.125 |
| 0.061 | 2.50 | 0.002 | 0.062 |
| 0.030 | 2.50 | 0.001 | 0.031 |
| 3.80 | 5 | 0.20 | 4 |
| 1.90 | 5 | 0.10 | 2 |
| 0.95 | 5 | 0.05 | 1 |

| | | | |
|---|---|---|---|
| 0.475 | 5 | 0.025 | 0.50 |
| 0.237 | 5 | 0.012 | 0.25 |
| 0.119 | 5 | 0.006 | 0.125 |
| 0.059 | 5 | 0.003 | 0.062 |
| 0.030 | 5 | 0.002 | 0.031 |
| 3.60 | 10 | 0.40 | 4 |
| 1.80 | 10 | 0.20 | 2 |
| 0.90 | 10 | 0.10 | 1 |
| 0.45 | 10 | 0.05 | 0.50 |
| 0.225 | 10 | 0.025 | 0.25 |
| 0.112 | 10 | 0.012 | 0.125 |
| 0.056 | 10 | 0.006 | 0.062 |
| 0.028 | 10 | 0.003 | 0.031 |
| 3.20 | 20 | 0.80 | 4 |
| 1.60 | 20 | 0.40 | 2 |
| 0.80 | 20 | 0.20 | 1 |
| 0.40 | 20 | 0.10 | 0.50 |
| 0.20 | 20 | 0.05 | 0.25 |
| 0.10 | 20 | 0.025 | 0.125 |
| 0.05 | 20 | 0.012 | 0.062 |
| 0.025 | 20 | 0.006 | 0.031 |
| 0 | 100 | 4 | 4 |
| 0 | 100 | 2 | 2 |
| 0 | 100 | 1 | 1 |
| 0 | 100 | 0.50 | 0.50 |
| 0 | 100 | 0.25 | 0.25 |
| 0 | 100 | 0.125 | 0.125 |
| 0 | 100 | 0.062 | 0.062 |
| 0 | 100 | 0.031 | 0.031 |

**Supplementary Table S2. Glucose-derived carbon concentrations used for species-specific growth assays.** Values are shown in g C $L^{-1}$ and were rounded for clarity.

| Glucose (g C $L^{-1}$) | g C $L^{-1}$ |
|---|---|
| 1 | 0.070 |
| 2 | 0.088 |
| 3 | 0.110 |
| 4 | 0.137 |
| 5 | 0.172 |
| 6 | 0.215 |
| 7 | 0.268 |
| 8 | 0.336 |
| 9 | 0.419 |
| 10 | 0.524 |
| 11 | 0.655 |
| 12 | 0.819 |
| 13 | 1.02 |
| 14 | 1.28 |
| 15 | 1.60 |
| 16 | 2 |

**Supplementary Table S3. Effects of resource gradients on community composition.** Permutational multivariate analysis of variance (PERMANOVA) testing the marginal effects of total carbon concentration and resource complexity on community composition. Analyses were performed on Bray-Curtis dissimilarities calculated from endpoint relative abundances (9'999 permutations). $R^2$ values indicate the proportion of total compositional variance explained by each term.

| Source of variation | Df | Sum of squares | $R^2$ | $F$ | $p$-value |
|---|---|---|---|---|---|
| Total carbon concentration | 1 | 90.36 | 0.546 | 968.9 | 0.0001 |
| Resource complexity (% R2A carbon contribution) | 1 | 3.68 | 0.022 | 39.5 | 0.0001 |
| Residual | 765 | 71.35 | 0.431 | — | — |
| Total | 767 | 165.39 | 1.000 | — | — |

**Supplementary Table S4. Effects of total carbon concentration and species identity on monoculture growth traits.** For each growth trait, linear models were fitted with standardized total carbon concentration (carbon), species identity, and their interaction as predictors. Shown are *F* statistics with numerator and denominator degrees of freedom and corresponding *p* values. Model fit is summarized using $R^2$ for additive (carbon + species) and interaction (carbon × species) models.

| Trait | Carbon effect | Species effect | Carbon × species | $R^2$ additive | $R^2$ interaction |
|---|---|---|---|---|---|
| Lag time | $F(1,44)=7.3$, $p=0.0098$ | $F(2,44)=0.86$, $p=0.43$ | $F(2,42)=9.4$, $p=4.2\times10^{-4}$ | 0.17 | 0.43 |
| Max growth rate | $F(1,44)=0.54$, $p=0.47$ | $F(2,44)=20$, $p=7.9\times10^{-7}$ | $F(2,42)=1.5$, $p=0.24$ | 0.48 | 0.51 |
| Carrying capacity | $F(1,44)=130$, p=$1.5\times10^{-14}$ | $F(2,44)=2.5$, $p=0.096$ | $F(2,42)=0.46$, $p=0.64$ | 0.75 | 0.76 |
| AUC | $F(1,44)=70$, p=$1.2\times10^{-10}$ | $F(2,44)=3.4$, $p=0.041$ | $F(2,42)=1.3$, $p=0.30$ | 0.64 | 0.66 |

**Supplementary Table S5. Pairwise species comparisons for maximum growth rate.** Tukey-adjusted pairwise contrasts based on estimated marginal means from linear models of maximum growth rate, averaged across the carbon gradient. Degrees of freedom correspond to the residual df of the fitted model. Negative estimates indicate lower maximum growth rate for the first species in the contrast.

| Contrast | Estimate | SE | df | *t* | *p* (Tukey) |
|---|---|---|---|---|---|
| *C. koseri* − *A. caviae* | $-1.44\times10^{-4}$ | $4.8\times10^{-5}$ | 42 | −2.98 | 0.013 |
| *C. koseri* − *P. chlororaphis* | $-3.06\times10^{-4}$ | $4.8\times10^{-5}$ | 42 | −6.34 | <0.001 |
| *A. caviae* − *P. chlororaphis* | $-1.62\times10^{-4}$ | $4.8\times10^{-5}$ | 42 | −3.36 | 0.0047 |

**Supplementary Table S6. Pairwise species comparisons for area under the growth curve (AUC).** Tukey-adjusted pairwise contrasts based on estimated marginal means from linear models of AUC, averaged across the carbon gradient. Degrees of freedom correspond to the residual df of the fitted model. Negative estimates indicate lower maximum growth rate for the first species in the contrast. No pairwise differences remained statistically significant after Tukey adjustment.

| Contrast | Estimate | SE | df | *t* | *p* (Tukey) |
|---|---|---|---|---|---|
| *C. koseri* − *A. caviae* | −0.06 | 0.65 | 42 | −0.09 | 0.995 |
| *C. koseri* − *P. chlororaphis* | −1.51 | 0.65 | 42 | −2.33 | 0.063 |
| *A. caviae* − *P. chlororaphis* | −1.45 | 0.65 | 42 | −2.23 | 0.077 |

**Supplementary Table S7. Cross-strain predictive models of community dominance.** Linear models testing whether monoculture growth traits predict relative abundance across strains. Analyses were performed separately for the full carbon gradient and the intermediate transition zone (0.25-1.0 g C $L^{-1}$). Unique $R^2$ indicates the additional variance explained by each trait after accounting for the others. Marginal $R^2$ indicates variance explained by each trait alone.

**Full carbon gradient**

$n$ = 144; Full model $R^2$ = 0.493; Adjusted $R^2$ = 0.475

| Trait | Standardized β | p-value | Unique $R^2$ | Marginal $R^2$ |
|---|---|---|---|---|
| Lag time | -0.255 | <0.001 | 0.314 | 0.161 |
| Maximum growth rate | 0.241 | <0.001 | 0.136 | 0.018 |
| AUC | 0.099 | 0.002 | 0.039 | 0.017 |

**Transition zone (0.25–1.0 g C $L^{-1}$)**

$n$ = 72; Full model $R^2$ = 0.730; Adjusted $R^2$ = 0.709

| Trait | Standardized β | *p*-value | Unique $R^2$ | Marginal $R^2$ |
|---|---|---|---|---|
| Lag time | -0.281 | <0.001 | 0.439 | 0.112 |
| Maximum growth rate | 0.383 | <0.001 | 0.349 | 0.083 |
| AUC | 0.045 | 0.063 | 0.015 | 0.029 |

**Supplementary Table S8. Linear model ANOVA for community biomass (OD).** Community biomass was modelled as a function of total carbon concentration, resource complexity (% R2A carbon contribution), and their interaction using a linear model. Predictors were z-standardised prior to analysis.

| Term | df | Sum Sq | Mean Sq | *F* value | *p*-value |
|---|---|---|---|---|---|
| Total carbon | 1 | 4.775 | 4.775 | 878.68 | <0.001 |
| Resource complexity | 1 | 0.282 | 0.282 | 51.84 | <0.001 |
| Carbon × complexity | 1 | 0.556 | 0.556 | 102.32 | <0.001 |
| Residuals | 764 | 4.152 | 0.005 | — | — |

**Supplementary Table S9. $R^2$ decomposition for community biomass.** Variance explained by total carbon, resource complexity, and their interaction was quantified by comparing $R^2$ values from nested linear models. Main effects represent unique contributions after accounting for the other predictor; interaction represents additional variance explained beyond additive effects.

| Component | $R^2$ |
|---|---|
| Total carbon (main effect) | 0.469 |
| Resource complexity (main) | 0.029 |
| Interaction | 0.057 |
| Total (additive model) | 0.515 |
| Total (interaction model) | 0.571 |

**Supplementary Table S10. Linear model ANOVA for Shannon diversity.** Shannon diversity was modelled as a function of total carbon concentration, resource complexity, and their interaction.

| Term | F value | $Df_1$ | $Df_2$ | *p*-value |
|---|---|---|---|---|
| Total carbon | 846.0 | 1 | 765 | <0.001 |
| Resource complexity | 156.0 | 1 | 765 | <0.001 |
| Carbon × complexity | 4.15 | 1 | 764 | 0.042 |

**Supplementary Table S11. $R^2$ contributions of total carbon, resource complexity, and their interaction to Shannon diversity.** Variance explained by each predictor was quantified by comparing $R^2$ values of nested linear models. Main effects represent unique variance attributable to each resource axis; interaction represents additional variance beyond additive effects.

| Component | $R^2$ |
|---|---|
| Total carbon (main effect) | 0.479 |
| Resource complexity (main) | 0.088 |
| Interaction | 0.002 |

**Supplementary Table S12.** Full primer sequences for PCR-1.

| Tagged fusion primers for PCR-1 | Sequence 5'-3' | Internal tag sequence |
|---|---|---|
| **iTru_A_16S_HAMBI_341F** | ACACTCTTTCCCTACACGACGCTCTTCCGATCT**GGTAC**CCTACGGGAGGCAGCAG | GGTAC |
| **iTru_B_16S_HAMBI_341F** | ACACTCTTTCCCTACACGACGCTCTTCCGATCT**cAACAC**CCTACGGGAGGCAGCAG | cAACAC |
| **iTru_C_16S_HAMBI_341F** | ACACTCTTTCCCTACACGACGCTCTTCCGATCT**atCGGTT**CCTACGGGAGGCAGCAG | atCGGTT |
| **iTru_D_16S_HAMBI_341F** | ACACTCTTTCCCTACACGACGCTCTTCCGATCT**tcgGTCAA**CCTACGGGAGGCAGCAG | tcgGTCAA |
| **iTru_E_16S_HAMBI_341F** | ACACTCTTTCCCTACACGACGCTCTTCCGATCT**AAGCG**CCTACGGGAGGCAGCAG | AAGCG |
| **iTru_F_16S_HAMBI_341F** | ACACTCTTTCCCTACACGACGCTCTTCCGATCT**gCCACA**CCTACGGGAGGCAGCAG | gCCACA |
| **iTru_G_16S_HAMBI_341F** | ACACTCTTTCCCTACACGACGCTCTTCCGATCT**ctGGATG**CCTACGGGAGGCAGCAG | ctGGATG |
| **iTru_H_16S_HAMBI_341F** | ACACTCTTTCCCTACACGACGCTCTTCCGATCT**tgaTTGAC**CCTACGGGAGGCAGCAG | tgaTTGAC |
| **iTru_1_16S_HAMBI_518R** | GTGACTGGAGTTCAGACGTGTGCTCTTCCGATCT**AGGAA**ATTACCGCGGCTGCTGG | AGGAA |
| **iTru_2_16S_HAMBI_518R** | GTGACTGGAGTTCAGACGTGTGCTCTTCCGATCT**gAGTGG**ATTACCGCGGCTGCTGG | gAGTGG |
| **iTru_3_16S_HAMBI_518R** | GTGACTGGAGTTCAGACGTGTGCTCTTCCGATCT**ccACGTC**ATTACCGCGGCTGCTGG | ccACGTC |
| **iTru_4_16S_HAMBI_518R** | GTGACTGGAGTTCAGACGTGTGCTCTTCCGATCT**ttcTCAGC**ATTACCGCGGCTGCTGG | ttcTCAGC |
| **iTru_5_16S_HAMBI_518R** | GTGACTGGAGTTCAGACGTGTGCTCTTCCGATCT**CTAGG**ATTACCGCGGCTGCTGG | CTAGG |
| **iTru_6_16S_HAMBI_518R** | GTGACTGGAGTTCAGACGTGTGCTCTTCCGATCT**tGCTTA**ATTACCGCGGCTGCTGG | tGCTTA |
| **iTru_7_16S_HAMBI_518R** | GTGACTGGAGTTCAGACGTGTGCTCTTCCGATCT**gcGAAGT**ATTACCGCGGCTGCTGG | gcGAAGT |
| **iTru_8_16S_HAMBI_518R** | GTGACTGGAGTTCAGACGTGTGCTCTTCCGATCT**aatCCTAT**ATTACCGCGGCTGCTGG | aatCCTAT |
| **iTru_9_16S_HAMBI_518R** | GTGACTGGAGTTCAGACGTGTGCTCTTCCGATCT**ATCTG**ATTACCGCGGCTGCTGG | ATCTG |
| **iTru_10_16S_HAMBI_518R** | GTGACTGGAGTTCAGACGTGTGCTCTTCCGATCT**gAGACT**ATTACCGCGGCTGCTGG | gAGACT |
| **iTru_11_16S_HAMBI_518R** | GTGACTGGAGTTCAGACGTGTGCTCTTCCGATCT**cgATTCC**ATTACCGCGGCTGCTGG | cgATTCC |

| | | |
|---|---|---|
| **iTru_12_16S_HAMBI_518R** | GTGACTGGAGTTCAGACGTGTGCTCTTCCGATCT**tctCAATC**ATTACCGCGGCTGCTGG | tctCAATC |

**Supplementary Table S13.** Full primer sequences for PCR-2.

| Name | Sequence 5'-3' | Index Label | Index seq |
|---|---|---|---|
| **iTru5_01_A** | AATGATACGGCGACCACCGAGATCTACACACCGACAAACACTCTTTCCCTAC | A | ACCGACAA |
| **iTru5_01_B** | AATGATACGGCGACCACCGAGATCTACACAGTGGCAAACACTCTTTCCCTAC | B | AGTGGCAA |
| **iTru5_01_C** | AATGATACGGCGACCACCGAGATCTACACCACAGACTACACTCTTTCCCTAC | C | CACAGACT |
| **iTru5_01_D** | AATGATACGGCGACCACCGAGATCTACACCGACACTTACACTCTTTCCCTAC | D | CGACACTT |
| **iTru5_01_E** | AATGATACGGCGACCACCGAGATCTACACGACTTGTGACACTCTTTCCCTAC | E | GACTTGTG |
| **iTru5_01_F** | AATGATACGGCGACCACCGAGATCTACACGTGAGACTACACTCTTTCCCTAC | F | GTGAGACT |
| **iTru5_01_G** | AATGATACGGCGACCACCGAGATCTACACGTTCCATGACACTCTTTCCCTAC | G | GTTCCATG |
| **iTru5_01_H** | AATGATACGGCGACCACCGAGATCTACACTAGCTGAGACACTCTTTCCCTAC | H | TAGCTGAG |
| **iTru7_101_01** | CAAGCAGAAGACGGCATACGAGATGGTAACGTGTGACTGGAGTTCAG | 1 | ACGTTACC |
| **iTru7_101_02** | CAAGCAGAAGACGGCATACGAGATCAACACAGGTGACTGGAGTTCAG | 2 | CTGTGTTG |
| **iTru7_101_03** | CAAGCAGAAGACGGCATACGAGATACACCTCAGTGACTGGAGTTCAG | 3 | TGAGGTGT |
| **iTru7_101_04** | CAAGCAGAAGACGGCATACGAGATCATGGATCGTGACTGGAGTTCAG | 4 | GATCCATG |
| **iTru7_101_05** | CAAGCAGAAGACGGCATACGAGATTGATAGGCGTGACTGGAGTTCAG | 5 | GCCTATCA |
| **iTru7_101_06** | CAAGCAGAAGACGGCATACGAGATCGGTTGTTGTGACTGGAGTTCAG | 6 | AACAACCG |
| **iTru7_101_07** | CAAGCAGAAGACGGCATACGAGATCAACGAGTGTGACTGGAGTTCAG | 7 | ACTCGTTG |
| **iTru7_101_08** | CAAGCAGAAGACGGCATACGAGATACCATAGGGTGACTGGAGTTCAG | 8 | CCTATGGT |

### Bibliography

1. Glenn, T. C., et al., ‘Adapterama II: Universal Amplicon Sequencing on Illumina Platforms (TaggiMatrix)’, *PeerJ*, 7 (2019), e7786, https://doi.org/10.7717/peerj.7786.

2. Rohland, N., and D. Reich, ‘Cost-Effective, High-Throughput DNA Sequencing Libraries for Multiplexed Target Capture’, *Genome Research*, 22/5 (2012), 939–46, https://doi.org/10.1101/gr.128124.111.
